# Aspect influences vegetation type in the Western Ghats

**DOI:** 10.1101/2025.07.23.666346

**Authors:** Devi Maheshwori, Shreyas Managave, Girish Jathar, Sham Davande

## Abstract

Western Ghats (WG) of India is one of the important biodiversity hotspots characterized by high levels of species endemism. The length of the rainy season and total rainfall are known to influence regional scale vegetation pattern. A recent study identified two aspect-related modes that influence landscape-scale tree cover (TC) and canopy height (CH) in the WG: north-facing slopes have a higher TC and CH than the south-facing slopes and (ii) higher TC and CH on the west-facing slopes than the east-facing slopes. However, it remains unclear whether these aspect-related asymmetries extend to the vegetation types. Here, with the help of seasonal NDVI variations, we demonstrate that aspect influence vegetation type as well. In general, north and west facing aspects support higher proportion of evergreen forests than the south and east facing slopes. In contrast, the dry deciduous and thorn forests relatively dominate the south and east-facing slopes. This natural heterogeneity in the vegetation type linked to aspect needs to considered while designing ecological research and management plans in the WG.

## 1. Introduction

Continental rifting and separation of India from the Seychelles which occurred during the late Cretaceous created a passive margin with an elevated area along the west coast of India (Gunnell and Harbor, 2008). The present-day remnant of which constitutes a sequence of hills running for about 1600 km parallel to the west coast of India and covering an area of ∼160000 km^2^ (Das et al., 2006), the Western Ghats (WG). The WG are one of the biodiversity hotspots having high species diversity and endemism (Myers et al., 2000). Although, the factors affecting regional vegetation distribution in the WG is known for a long (Champion and Seth 1968; Pascal 1988; Gadgil, 1996), those influencing vegetation within a sanctuary, important for understanding spatial variability of species richness and for conserving endemic flora and fauna, is not fully understood.

Evergreen, semi-evergreen, moist deciduous and dry deciduous are the main types of forests in the WG (Champion and Seth, 1968). The length of the rainy season and total rainfall control the regional vegetation distribution in the WG (Champion and Seth, 1968; Pascal 1988; Gadgil, 1996; Gimaret-Carpentier et al., 2003; Nagendra and Ghate, 2003; Prasad et al., 2008). Climatic variables are the most important factors controlling variation in woody species composition at regional and landscape scales (Page and Shanker, 2020). Two regional scale patterns have been identified to control the vegetation distribution in the WG. The south to north increase in the length of the dry season has resulted in lesser vegetation vigor northwards (Nagendra and Ghate, 2003). The drier northern part is associated with lower plant diversity than the southern part (Nagendra and Ghate, 2003). Overall, the species richness of woody plants increases monotonically from higher to lower latitudes with the species in the NWG exhibiting greater tolerance to temperature and rainfall seasonality (Page and Shanker, 2020). Going from east to west of the WG, the forest type changes from dry deciduous to moist deciduous to evergreen along the increasing rainfall gradient (Pascal 1988; Gadgil, 1996).

The role of aspect, the direction a given slope faces, in controlling vegetation type in the WG is realized mainly in terms of its relationship with the orographic precipitation (Champion and Seth, 1935; Gadgil, 1996). The WG acts as a barrier for moisture laden southwesterly winds, ensuring higher precipitation on windward facing western aspect than the leeward facing eastern aspect. This results in west to east gradient in the vegetation type: evergreen forest on west facing slopes which changes to moist deciduous to dry deciduous on eastward slopes (Gadgil, 1996).

The south-facing slopes in the northern hemispheres receive relatively stronger solar radiation than the north-facing slopes; the asymmetry in the solar radiation received increases poleward and with the slope angle at a given latitude (Smith and Bookhagen, 2021). As a result, the south-facing slopes in the northern Hemisphere have a relatively warmer, drier microclimate and soil with lower moisture content than the north-facing slopes (Geroy et al., 2011; Gutiérrez-Jurado et al., 2013; Zhou et al., 2013). This coupled with soil edaphic factors, leads to the north-facing slopes having a higher vegetation cover than the South-facing slopes in the Northern Hemisphere (Smith and Bookhagen, 2021; Maheshwori et al., 2025). The north-south (N-S) asymmetry in the vegetation cover thus established is common to mid-latitude regions (e.g. Champion and Seth 1935; Holland and Steyn, 1975; Badano et al., 2005, Singh, 2018). The influence of slope-aspect on vegetation distribution is recognized relatively late in the WG (Maheshwori et al., 2025).

The seasonal rainfall and asymmetric solar heating enable the slope-aspect to influence the landscape-scale tree cover and canopy height in the WG (Maheshwori et al., 2025). The aspect-related asymmetry in the vegetation structure is reflected in two modes: the North-aspect has higher tree cover (TC) and canopy height (CH) than the South-aspect; the West-aspect has higher TC and CH than the East-aspect. It is, however, not clear whether these asymmetries are also reflected in the vegetation type. Here, with the help of seasonal NDVI variations of vegetation, we test whether the aspect influences the landscape-scale vegetation type in the WG.

## 2. Materials and Methods

### 2.1. Regional Climate

The WG acts as an orographic barrier to the south-westerly monsoonal winds during southwest monsoon (SWM) season (June to September) which results in higher and lower rainfall on the windward (western margin) and leeward (eastern margin) sides of the WG, respectively (Gunnell, 1997) (Fig. 1). Only the southern part of WG receives additional rains during the northeast monsoon (NEM) season (October to December) (Fig. 1). This, in conjunction with an earlier onset and late withdrawal of SWM in the southern part of the WG, leads to a longer rainy season (∼260 days) in the southern part as compared to than in the northern part (∼130 days) (Fig. 1) (Singh 1986). The extreme southern part, however, receives less number of rainy days and lower rainfall (Guhathakurta et al., 2020).

**Figure 1.**
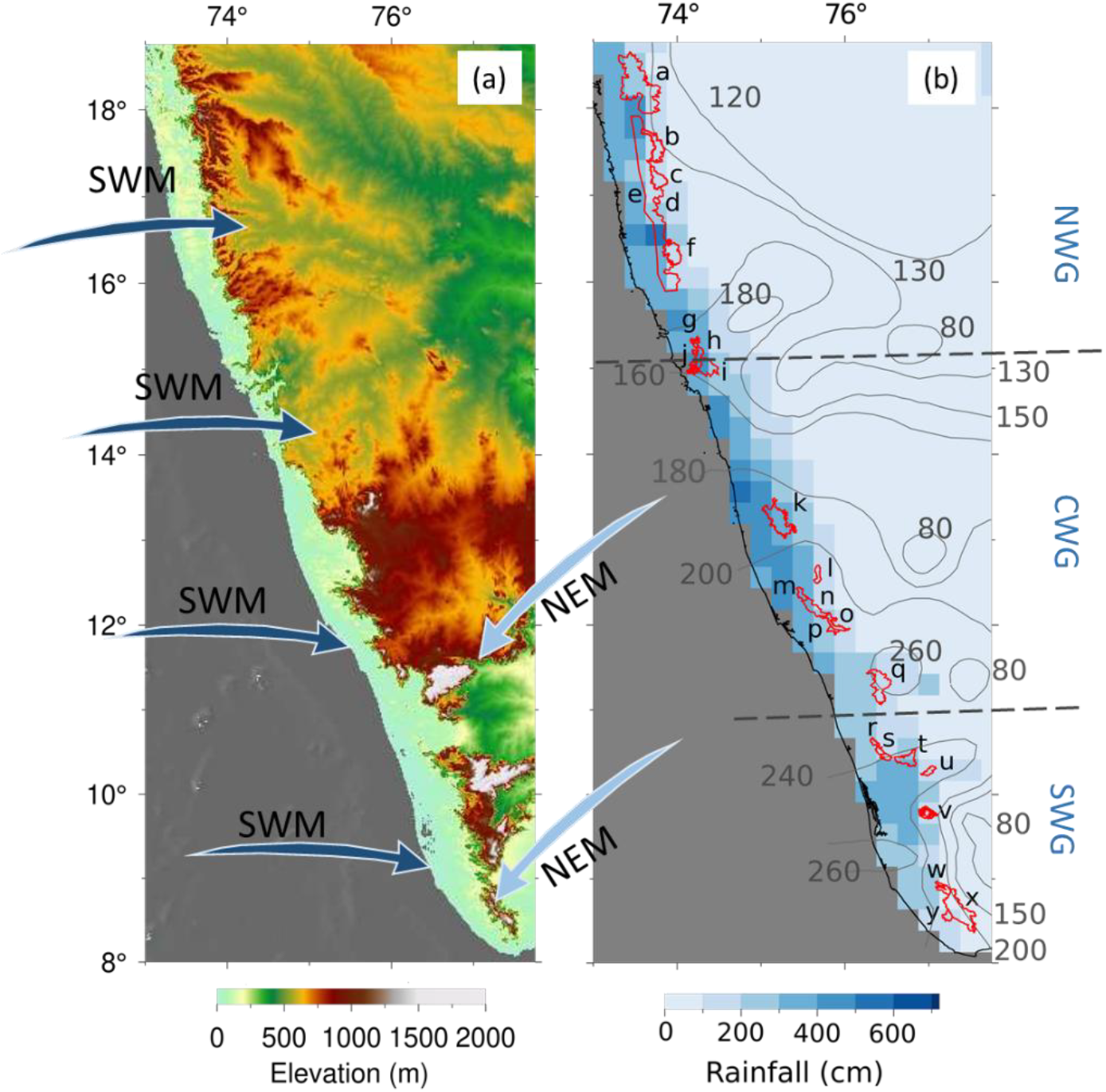
Altitudinal (based on ASTER Global Digital Elevation Model (a), rainfall (Pai et al., 2014) (b) and the length of rainy season (Singh., 1986) (b) distribution in the WG. The arrows in (a) indicate low level wind direction during the southwest (SWM) and northeast (NEM) monsoons. The protected areas (PAs) whose NDVI variations are studied in this work are highlighted by different colours and marked by letters ‘a’ to ‘y’ as: a-Tamhini-Lavasa-Raigad; b-Koyna; c-Chandoli; d-Vishalgad; e-West of the WG Escarpment; f-Radhangiri; g-Bhagwan Mahavir; h-Netravali; i-Anshi; j-Cotigao; k-Kudremukh; l-Pushpagiri; m-Talakaveri; n-New Brahmagiri; o-Brahmagiri; p-Aralam; q-New Amarambalam-Silent Valley; r-Peechi; s-Chimmony; t-Parambikulun; u-Indira Gandhi; v-Idukki; w-Shendurney; x-Kallakad Mundanthurai; y-Neyyar. NWG, CWG and SWG indicate northern, central and southern WG, respectively.

### 2.2. Methodology

The WG landscape is affected by anthropogenic activities (Rao et al., 2013). To minimize their effects of the aspect-related asymmetry in vegetation type, only protected areas (PAs) were considered in this study (Fig. 1). The same PAs (N = 25) that were considered by Maheshwori et al., (2025) were selected in this study. The boundaries of selected PAs were from the Ministry of Environment, Forest and Climate Change, Government of India’s Decision Support System (MoEF & CC-DSS). The boundaries of some of the PAs (Tamhini-Lavasa-Raigad, West of the WG Escarpment, New Brahmagiri, New Amarambalam-Silent Valley and Vishalgad) were drawn manually. Areas under plantations, human settlements and water bodies, deciphered through Google Earth images, were not considered.

To investigate the vegetation heterogeneity, Normalized Difference Vegetation Index (NDVI), derived from Sentinel-2 MSI Level-2A surface reflectance data at a 30 m resolution, was used. Data was processed in Google Earth Engine (GEE) and covered the period from October 2023 to May 2024 (monthly). NDVI values were calculated using reflectance in the near infra-red (NIR) and red (Red) region using the following formula:

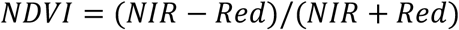

The wet season NDVI was derived from post-monsoon data with cloud cover below 20%, while the dry season NDVI represented the minimum NDVI value recorded during non-wet months.

Topographic variables including aspect, slope, and elevation, were computed from the 30 m resolution SRTM DEM (NASA JPL 2013). Aspect was categorized into eight cardinal directions: North (N), Northeast (NE), East (E), Southeast (SE), South (S), Southwest (SW), West (W) and Northwest (NW). The NDVI and topographic data were spatially integrated using Q-GIS (QGIS.org, 2024). Areas with missing NDVI values were excluded prior to converting the dataset into a gridded format containing latitude, longitude, and NDVI values. NDVI values were assigned to each latitude-longitude coordinate using the nearest neighbor interpolation (Franco-Lopez et al., 2001).

### 2.3. NDVI-based vegetation classes: a new approach to identify forest type

The monthly NDVI values of forests in the WG follows the maximum temperature and rainfall, and vary depending upon the vegetation type (Prabakaran et al., 2013; Ramachandra et al., 2016; Singh et al., 2020). In the WG, the minimum NDVI values occur during the dry summer season (i. e. April to June) whereas the maximum values occur during October and November (Ramachandra et al., 2016). Evergreen forests have high NDVI values during the wet as well as dry seasons (Singh et al., 2020). The moist deciduous forest has NDVI values similar to the evergreen forest during wet season but lower values during the dry season (Prasad et al., 2005; Singh et al., 2020). In comparison to the evergreen forests, the dry deciduous forests exhibit lower NDVI values during both the wet and dry seasons. The mean NDVI values (of February, November and April) decrease from moist deciduous to dry deciduous to scrub forests (Bawa et al., 2002). The lower NDVI values of dry deciduous forest could also be in part due to exposed bare ground during the dry season.

The seasonal and vegetation-type dependent variation in the NDVI values formed the basis for defining various vegetation classes (VGC) (Fig. 2). We divided the space of NDVI values during dry and wet seasons into 16 parts (representing 16 VGCs) (Fig. 2). The NDVI values of wet and dry period represent the NDVI values immediately after the cessation of the rainy season (i.e. of the first cloud-free image) and the lowest value during the non-rainy months, respectively. As the month which shows the lowest NDVI value varies inter-annually due to precipitation and temperature variability (Ramachandra et al., 2016), we chose the lowest NDVI value during the driest month from the non-rainy months (generally January to May), instead of the NDVI value of a particular month. Large number of pixels lie in the area represented by VGC-10 and VGC-11 (shown subsequently). To assess whether these pixels show any preference for particular aspect, we further divided VGC-10 and VGC-11 into six subclasses as shown in the Figure 2.

**Figure 2.**
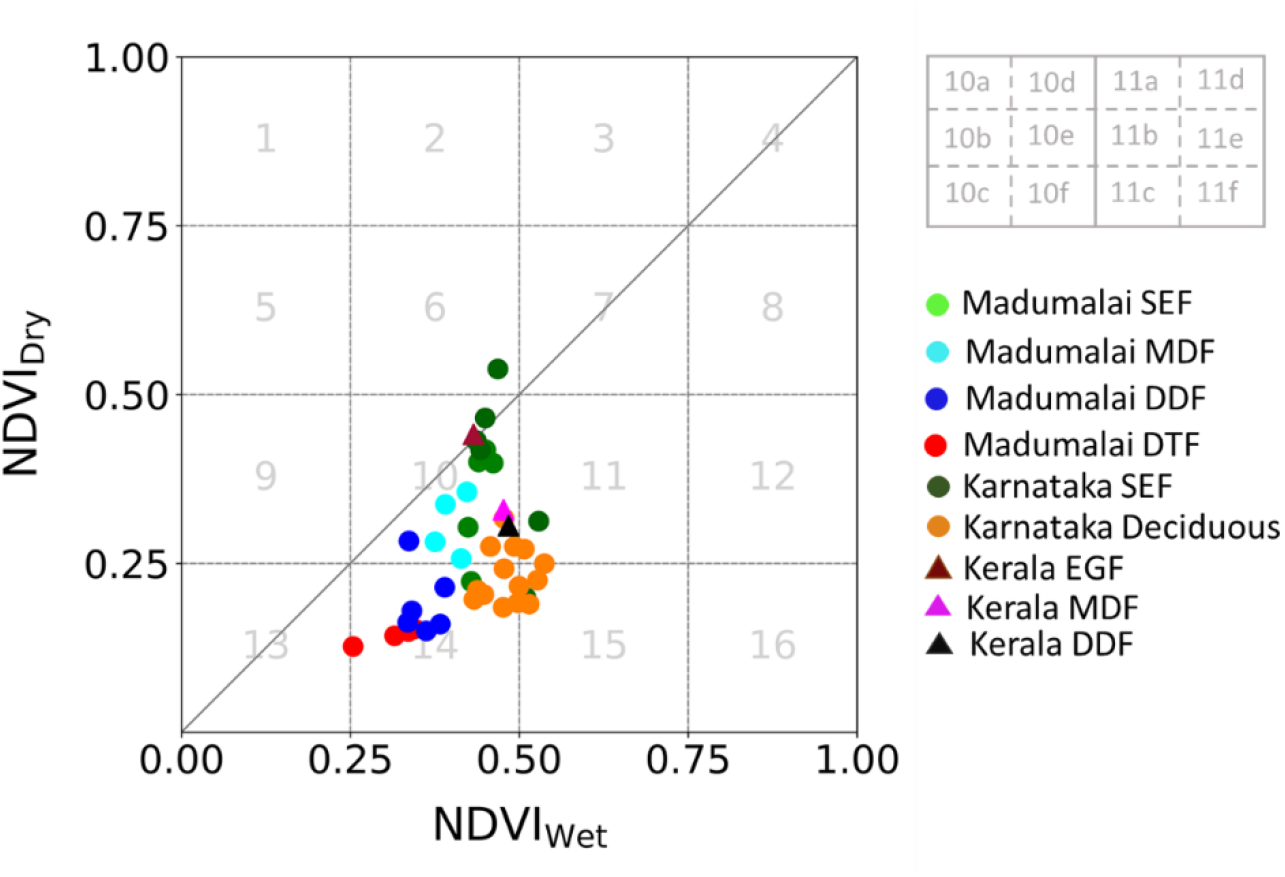
Vegetation classification scheme using NDVI values of the wet (after cessation of rains) and dry months. The numbers 1 to 16 indicate vegetation classes (VGC). The VGC-10 and VGC-11 were further divided into six parts as shown in the top right hand side of the figure. The data points indicate NDVI values of permanent long-term monitoring plots (Madumalai: Dattaraja et al., 2018; Karnataka: Sharma et al., 2021; Kerala: Sreejith et al., 2015). EGF, SEF, MDF, DDF and DTF are acronyms for evergreen, semi-evergreen, moist deciduous, dry deciduous and dry thorn forests, respectively.

Even though the forest types are broadly classified as evergreen, semi-evergreen, moist deciduous, dry deciduous etc., there lies a continuum between them as vegetation composition gradually grades between two coexisting forests types. Further, each forest type is not monotonous with respect to its vegetation composition but has various sub-types within them (Champion and Seth, 1968; Pascal et al., 2004). Even a subtype of a particular forest type could further show floristic heterogeneity. For examples, wet evergreen forests of southern Western Ghats were classified into 5 sub-types each having its own floristic composition (Pascal et al., 2004).

Most of the NDVI data points fall under VGC-7, VGC-10 (VGC-10d, VGC-e and VGC-f), VGC-11 (VGC-11a, VGC-11b and VGC-11c), VGC-14 and VGC15. Keeping the points mentioned in the previous paragraphs in mind, we propose that VGC -7 is likely to be dominated by evergreen species perhaps belonging to wet evergreen forests. VGC-14 likely to have domination of grass and deciduous trees belonging to dry deciduous and dry thorn forests. VGC-15 likely represents tree dominated deciduous species. VGC-10, the most dominant in the WG, is likely show gradation of dominant vegetation type from the top to bottom in VGC-10 region: evergreen/semi-evergreen at the top (VGC-10d) followed by moist deciduous in the middle (VGC-10e) and dry deciduous at the bottom (VGC-10f). VGC-11 at the top (VGC11-a) is more like VGC-7 and at bottom (VGC-11c) is more like VGC-15.

The assignments of VGCs and vegetation types were broadly corroborated by the NDVI values of permanent long-term monitoring plots whose forest type was verified by field-based observations (Fig. 2). However, it should be noted that these plots sample only a part of all the VGCs present in the WG (shown subsequently).

## 3. Results and Discussion

### 3.1 Median NDVI values on different aspects in PAs during dry period

The median NDVI values during the dry period were not the same on various aspects but showed a pattern wherein the N-, NW- and W-aspects in general showed higher NDVI values than the S-, SE- and E-aspects (Fig. 3). This suggested a higher proportion of deciduous species cover S-, SE- and E-aspects.

**Figure 3.**
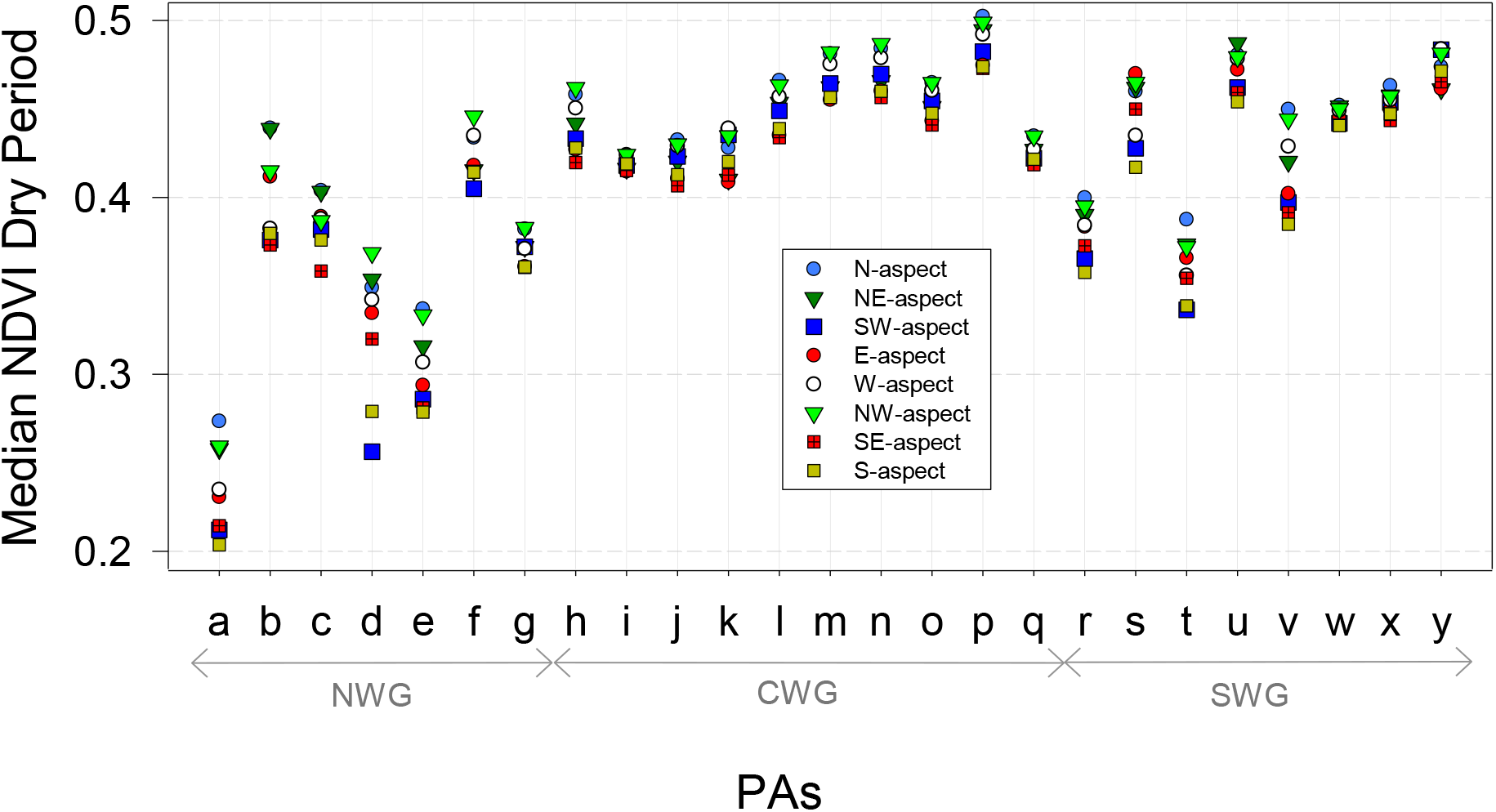
The median NDVI values during dry period on various aspects in different PAs. The abbreviations used are as in Figure. 1. NWG, CWG and SWG indicate northern, central and southern Western Ghats, respectively.

### 3.2 Asymmetry of vegetation types on N-, S-, W- and E-aspects

The N- and W-aspects showed a higher preference for VGC-7 (∼evergreen dominance) than the S- and E-aspects (Fig. 4 a, d). Contrarily, S- and E-aspect show higher preference for VGC-10 (Fig. 4 b, e) and VGC-14 (Fig. 4 c, f). The difference between the area covered by VGC-7, VGC-10 and VGC-14 on these aspects is shown in Table 1. The sub-classes VGC-10d, VGC-10e and VGC-10f also showed a higher preference for S- and E-aspects than for N- and W-aspects (Fig. 5). The preference of VGC-11a and VGC-11b was broadly similar to that of VGC-7; they showed higher abundance on the N- and W-aspect (Fig. 6). In the case of VGC-11c, most PAs showed subdued preference for aspect; the ones which has a higher abundance of VGC11-c showed preference for S- and W-aspects.

**Table 1.** Difference in the percent area covered by VGCs on N- and S-aspects and on W- and E-aspects.

|  | Vegetation Classes (VGC) |  |  |
| --- | --- | --- | --- |
|  | VGC-7 | VGC-10 | VGC-14 |
| N-aspect – S-aspect | $9 \pm 6$ | $-10 \pm 9$ | $-5 \pm 5$ |
| W-aspect – E-aspect | $7 \pm 6$ | $-10 \pm 8$ | $-2 \pm 3$ |

**Figure 4.**
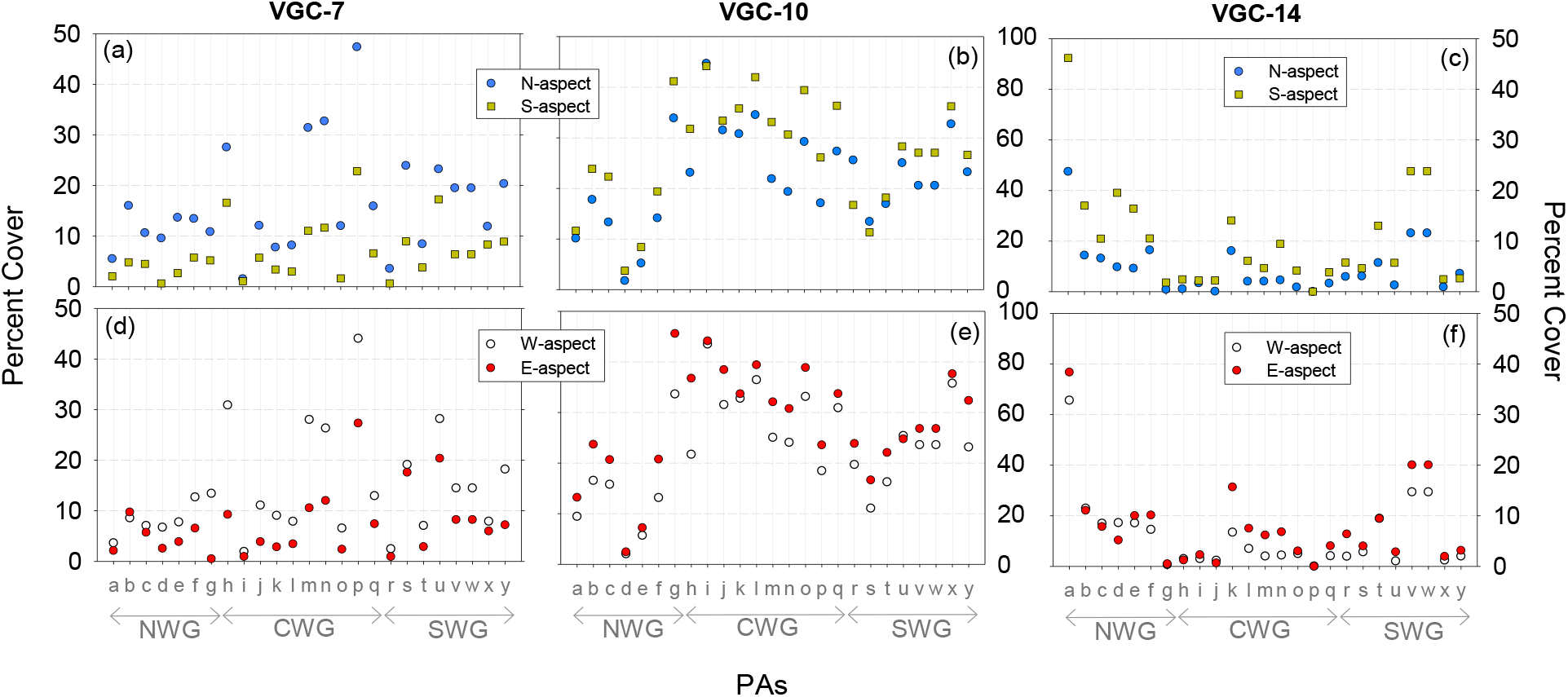
Percent area occupied by VGC-7 (left panels), VGC-10 (middle panels) and VGC-14 (right panels) on the N-aspect and S-aspect (top panels) and the W-aspect and E-aspect (bottom panels) in the studied PAs. For example, in PA ‘p’, VGC-7 occupied ∼47 % and ∼23 % of area covered by N-aspect and S-aspects, respectively. The abbreviations (‘a’ to ‘y’) used for the PAs are as in Figure. 1.

**Figure 5.**
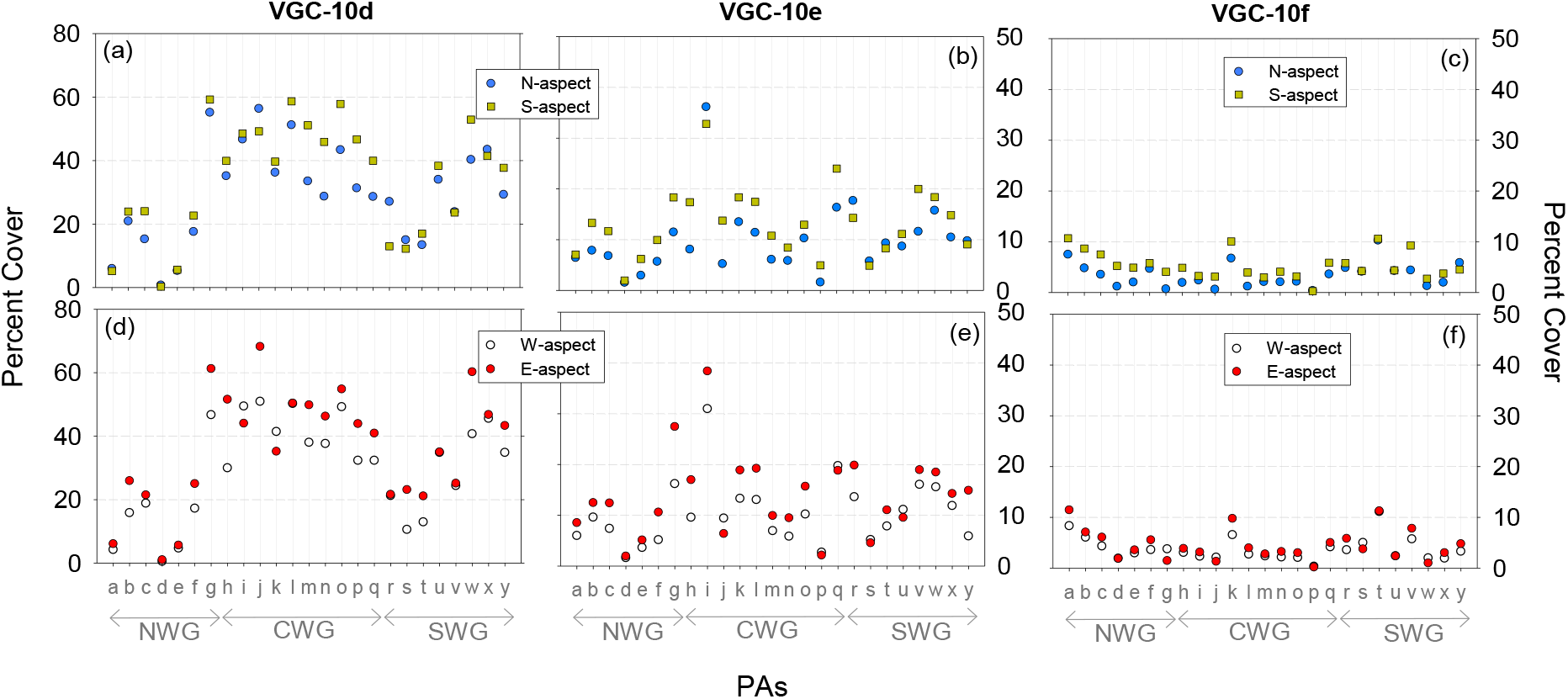
Percent area occupied by VGC-7 (left panels), VGC-10 (middle panels) and VGC-14 (right panels) on the N-aspect and S-aspect (top panels) and the W-aspect and E-aspect (bottom panels) in the studied PAs. The abbreviations (‘a’ to ‘y’) used for the PAs are as in Figure. 1.

**Figure 6.**
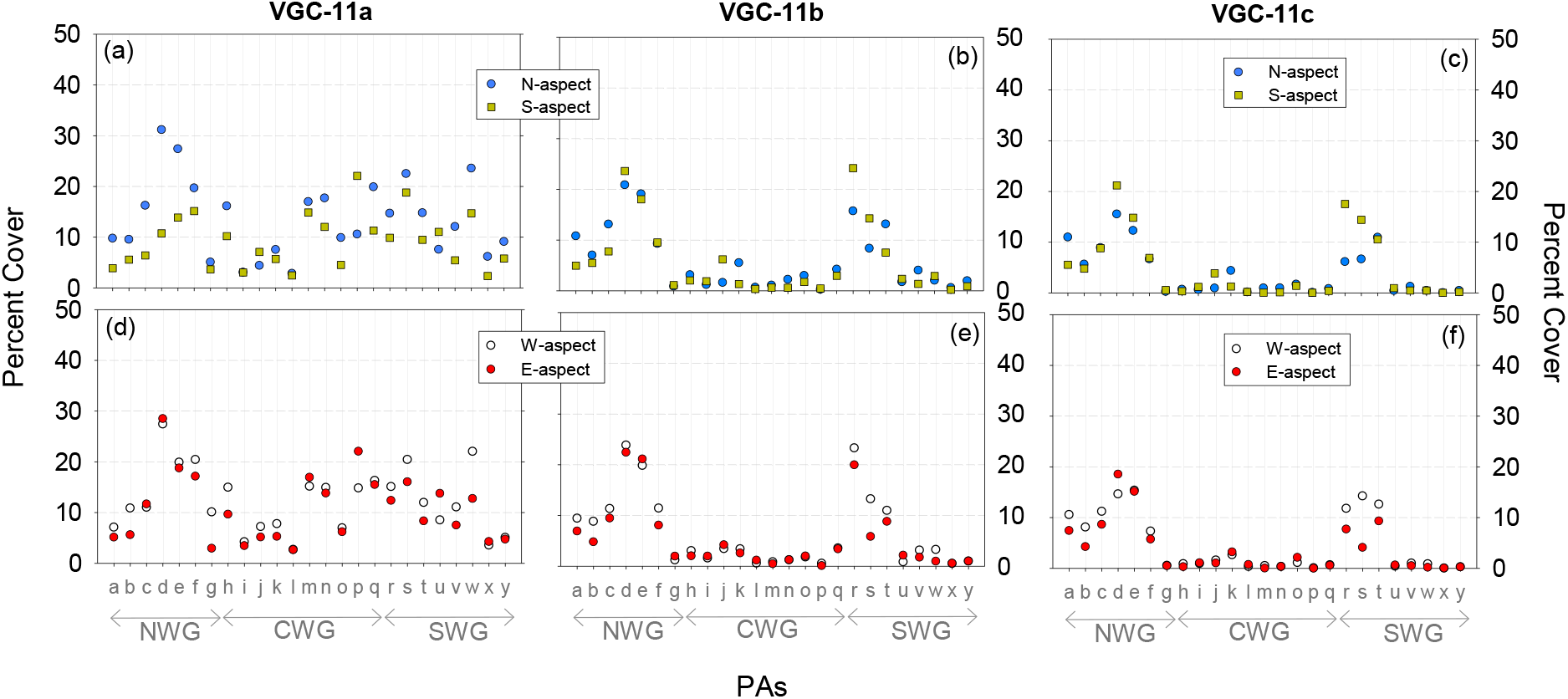
Percent area occupied by VGC-7 (left panels), VGC-10 (middle panels) and VGC-14 (right panels) on the N-aspect and S-aspect (top panels) and the W-aspect and E-aspect (bottom panels) in the studied PAs. The abbreviations (‘a’ to ‘y’) used for the PAs are as in Figure. 1

As the conditions becomes more conducive for the establishment of VGC-7, it expanded greater on the N-aspect than on the S-aspect; the slope 1.68 of a linear regression between percent area occupied by VGC-7 on N-aspect and S-aspect revealed this (Fig. 7a). Conversely, increasing conduciveness for the establishment of VGC-14 resulted in greater coverage of VGC-14 on the S-aspect compared to the N-aspect, as reflected by a regression slope of 0.48 between the percent area covered by VGC-14 on the N- and S-aspects (Fig. 7b). Similar patterns of variations were observed in the case VGC-7 (and VGC-14) on the W- and E-aspects.

**Figure 7.**
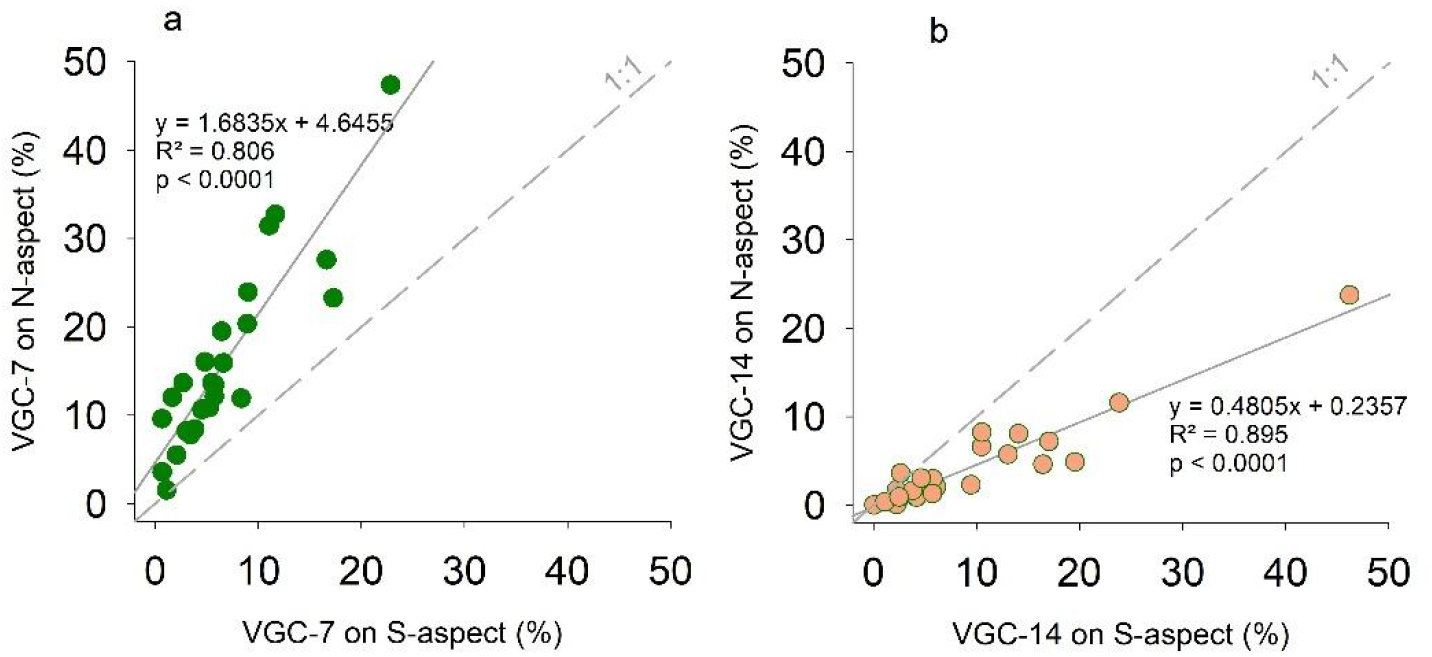
Percent area occupied by VGC-7 (left), VGC-14 (right) on the N-aspect and S-aspect.

This N-aspect to S-aspect and W-aspect to E-aspect trends in VGC-7 (wet conditions) and VGC-14 (dry conditions) observed here are consistent with the interpretations by Maheshwori et al., (2025) about the landscape scale distribution of TC and CH in the WG. Maheshwori et al., (2025) demonstrated the patterns of solar declination vis-à-vis monsoonal precipitation and orographic precipitation results in two modes of variability: i) the N-aspect support denser TC and taller CH than the S-aspect and ii) the W-aspect has a denser TC and taller CH than the E-aspect. Relatively higher heating of S-aspect post-rainy season (∼more soil water evaporation) and lower precipitation on E-aspect, possibly exacerbated by its (E-aspect’s) higher heating in the pre-noon hours, contribute to S- and E-aspects having higher VGC-14 on them. In contrast, lesser heating of N-aspect and higher precipitation on the W- aspect results in them having higher amount of VGC-7.

### 3.3. Vegetation composition of different aspects in the WG

Distribution of VGCs on all aspects (Fig. 8, 9, Supplementary Figures S1-S25) revealed that N-, NW- and W-aspects showed higher and lower preference, respectively, for VGC-7 and VGC-14. The highest and the lowest abundances (% covered) of VGC-7 was observed on NW- and SE-aspects, respectively. Further, VGC-10 was relatively more abundant of S-, SE- and E-aspect ands than on N-, NW-, W-aspects.

**Figure 8.**
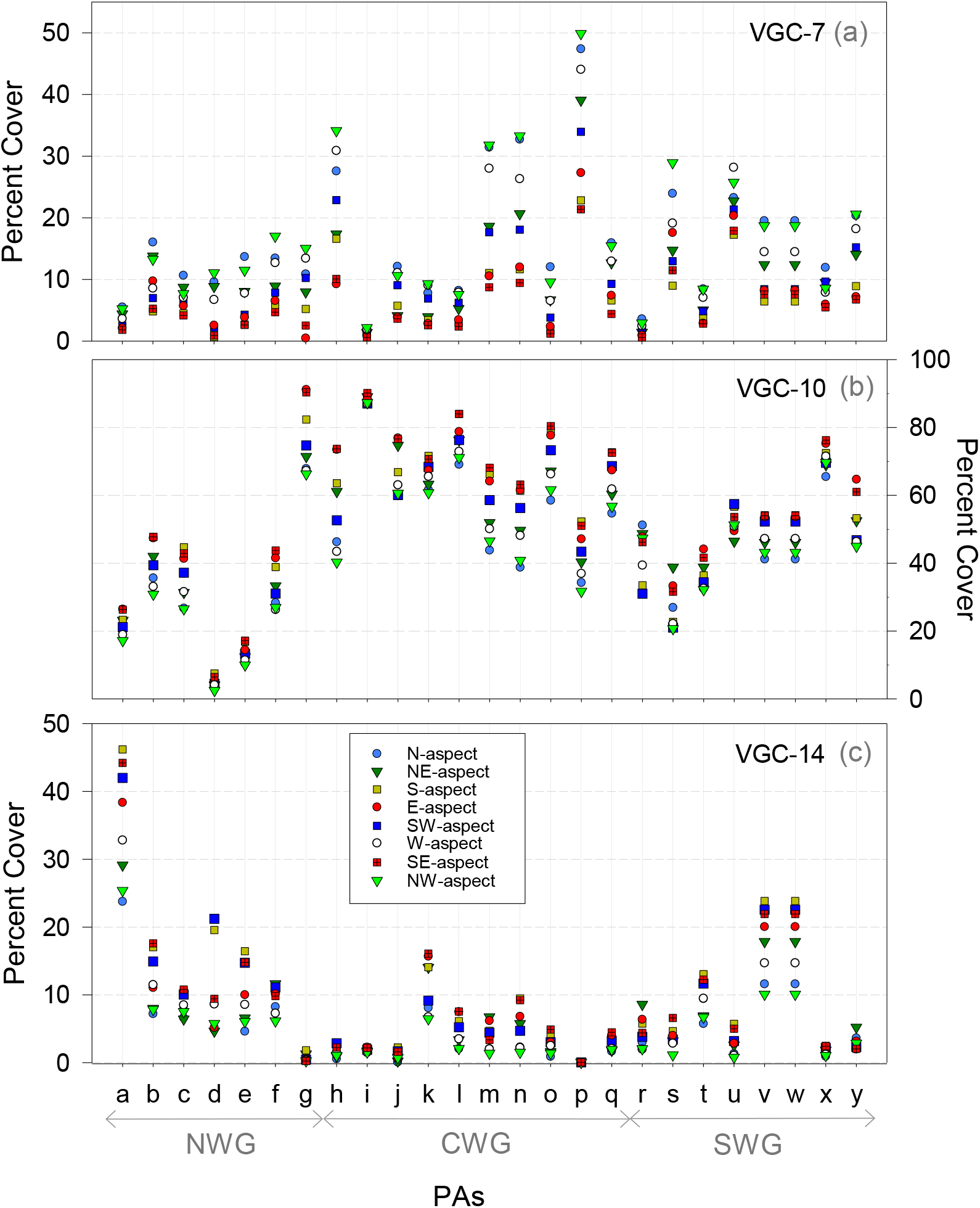
Percent area occupied by VGC-7 (top), VGC-10 (middle) and VGC-14 (bottom) on different aspect categories in the studied PAs. The legend shown in the bottom panel also holds for top and middle panels. The abbreviations used for the PAs are as in Figure. 1. NWG, CWG and SWG indicate northern, central and southern Western Ghats, respectively.

**Figure 9.**
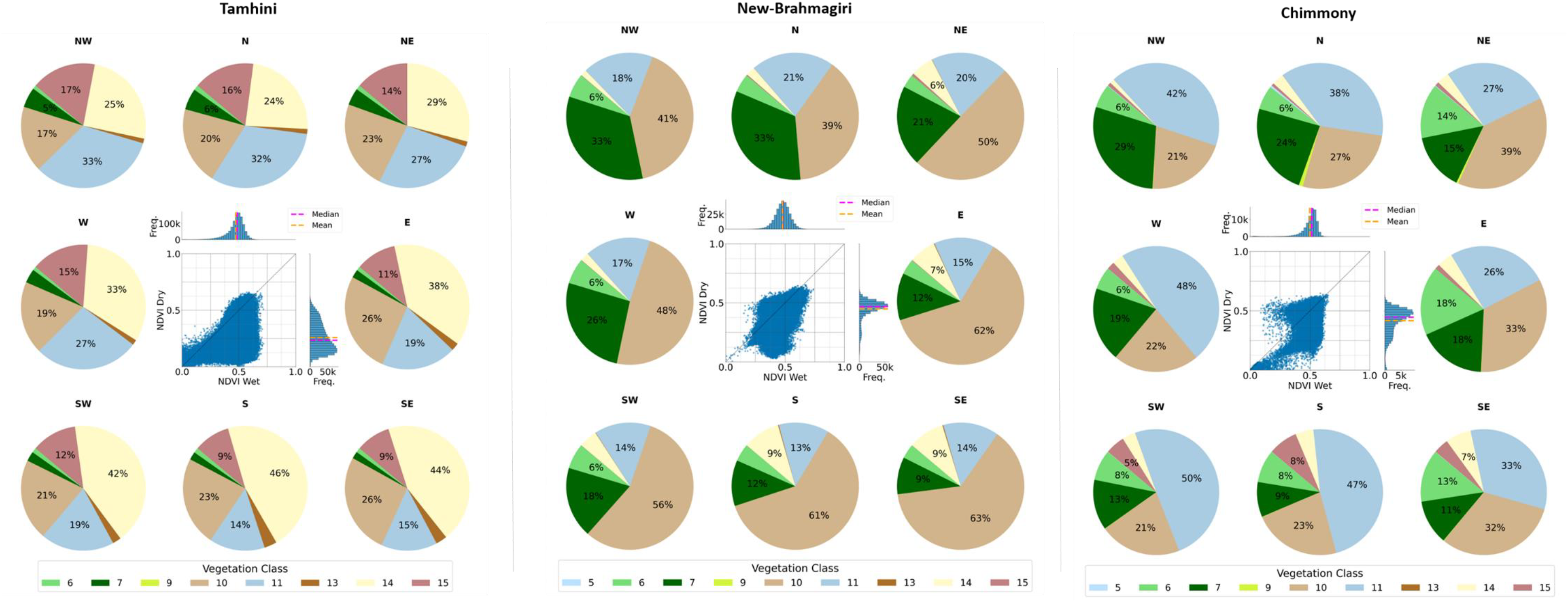
Pie charts showing area covered by different VGCs on 8 aspects in Tamhini in the NWG (left), New Brahmgiri in the CWG (middle) and Chimmony in the SWG (right). The scatter diagram in the middle shows NDVI values of vegetation in 30 × 30 m pixels during wet and dry seasons; the associated histograms show their frequency distribution. NW, N, NE, E, SE, S, SW and W represent northwest, north, northeast, east, southeast, south, southwest and west directions, respectively.

This study demonstrates that aspect, in addition to affecting tree cover and canopy height distribution (Maheshwori et al., 2025), also influence vegetation types in the WG. Two modes dominate: e.g., decreasing trend in VGC-7 from N-aspect to S-aspect and from W-aspect to E-aspect. These modes interact at the landscape scale and leads to a higher proportion of evergreen dominated forest (VGC-7) on N-, NW- and W-aspects and its lower proportion on S-, SE- and E-aspects; an observation similar to that reported for tree cover and canopy height (Maheshwori et al., 2025). This further implies that N-, NW- and W-aspects do not have ‘pristine forest’ and S-, SE- and E-aspects, ‘degraded forest’; it is a part of the natural spatial variability of vegetation type at the landscape scale.

## 5. Conclusions

The role of aspect in controlling vegetation type is well recognized in the mid-latitudes regions. Its influence in the WG has been realized mainly as the effect of orographic precipitation wherein the western side of the WG (i.e. the area lying west of the crest line of the WG hills) received higher precipitation than the eastern side. In this study we demonstrate that the vegetation type is also controlled by a N-S mode which results in the N-aspect having higher evergreen forest than the S-aspect. The combined effect of these two modes creates heterogeneous vegetation growth conditions on various aspects with the NW- and SE-aspects getting advantage (higher rainfall and reduced solar heating) and disadvantage (lower rainfall and increased solar heating) of these modes, respectively. This results in NW- and SE-facing slopes have the highest and lowest proportion of evergreen forests, respectively. This pattern is part of a natural system likely governed by interaction between the solar declination vis-àvis monsoonal rainfall and orographic precipitation. The aspect thus plays a crucial role in controlling vegetation structure (tree cover, canopy height and forest types) at the landscape scale in the WG; its influence should be considered while assessing landscape-scale biodiversity patterns and formulating effective forest management plans.

## Author contributions

Conceptualization: Shreyas Managave

Methodology: Shreyas Managave, Devi Maheshwori, Girish Jathar, Sham Davande

Formal Analysis: Devi Maheshwori, Shreyas Managave, Girish Jathar, Sham Davande

Investigation: Devi Maheshwori, Shreyas Managave, Girish Jathar, Sham Davande

Resources: Shreyas Managave

Writing—Original Draft: Shreyas Managave, Devi Maheshwori

Writing—Review and Editing: Shreyas Managave, Devi Maheshwori, Girish Jathar, Sham Davande

Supervision: Shreyas Managave

Funding Acquisition: Shreyas Managave

## Data Availability

Data will be made available upon request to the corresponding author.

