## Supplementary Figures for "Aspect influences vegetation type in the Western Ghats"

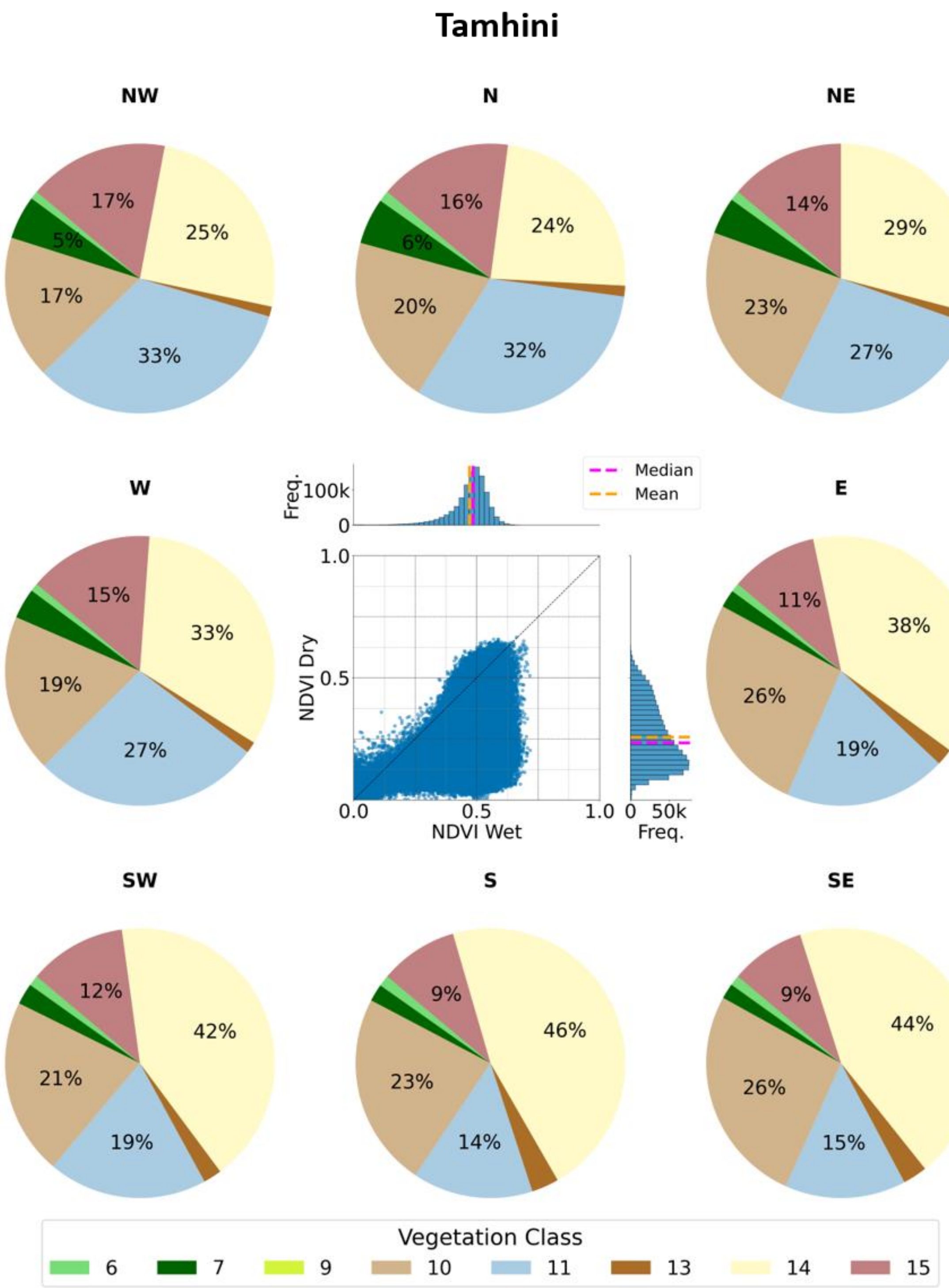


*Supplementary Figure S1. Pie charts showing area covered by different VGCs on 8 aspects in Tamhini The scatter diagram in the middle shows NDVI values of vegetation in 30 x 30 m pixels during wet and dry seasons; the associated histograms show their frequency distribution. NW, N, NE, E, SE, S, SW and W represent northwest, north, northeast, east, southeast, south, southwest and west directions, respectively.*

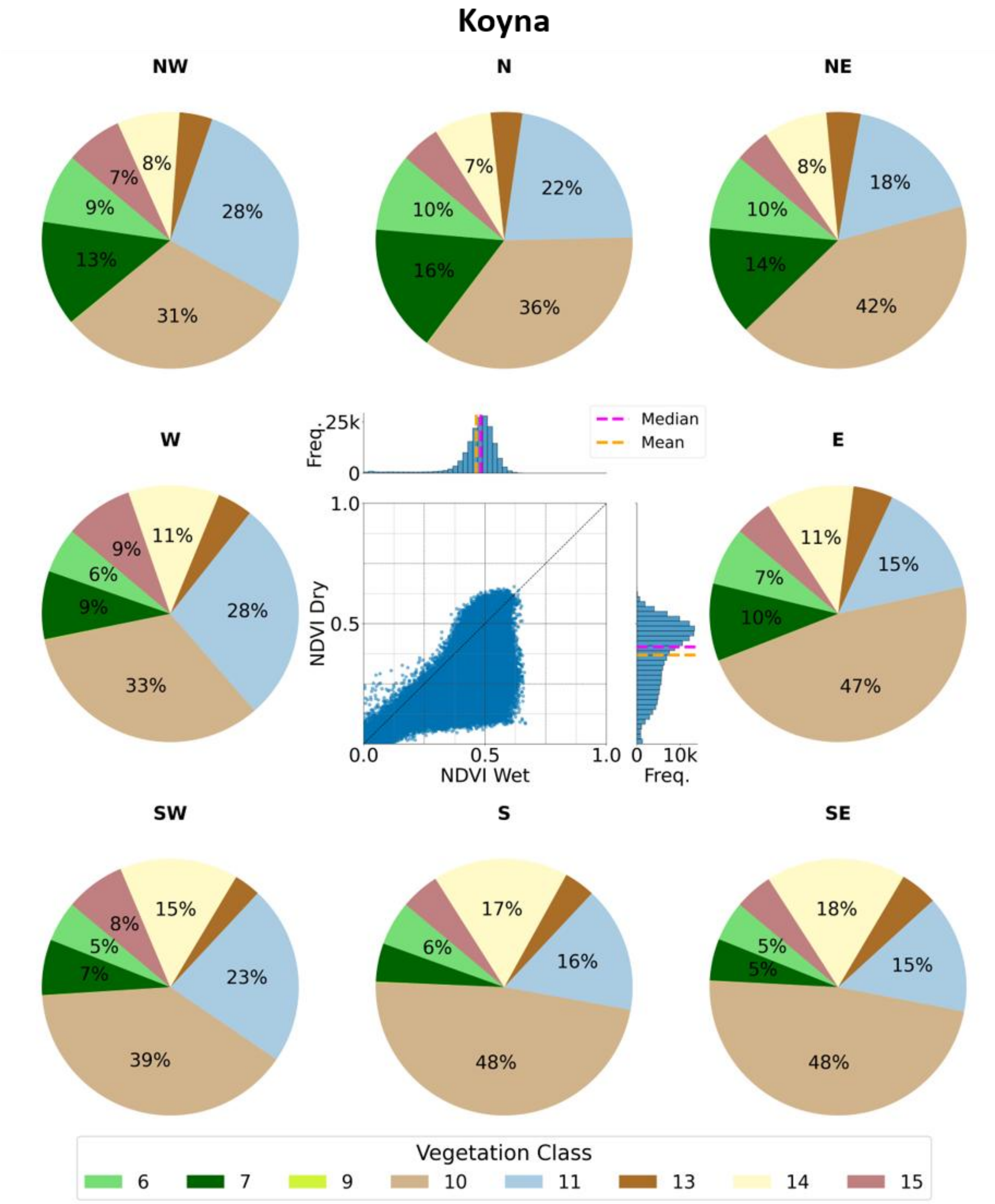


*Supplementary Figure S2. The same as Supplementary Figure S1 but for Koyna.*

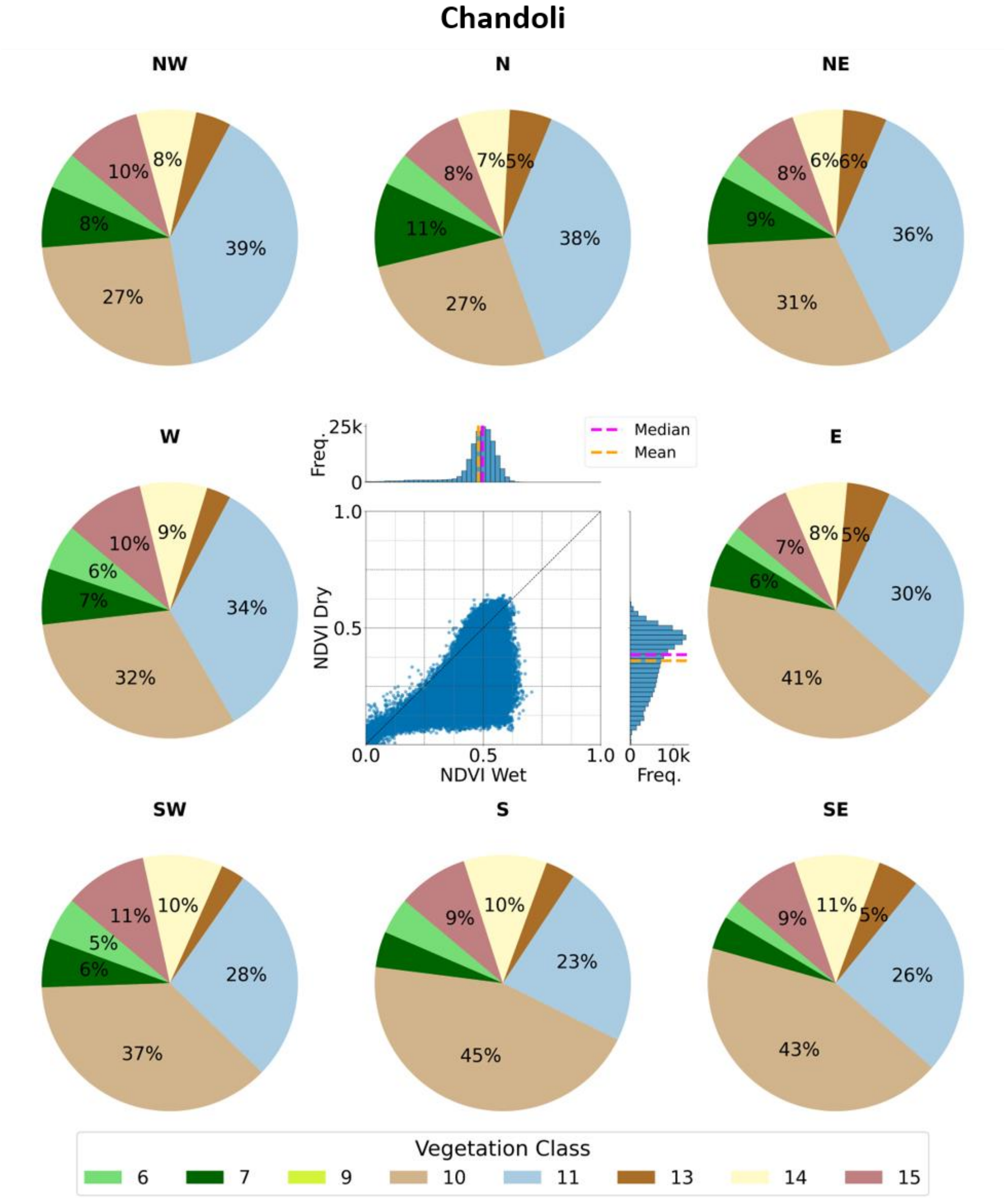


*Supplementary Figure S3. The same as Supplementary Figure S1 but for Chandoli.*

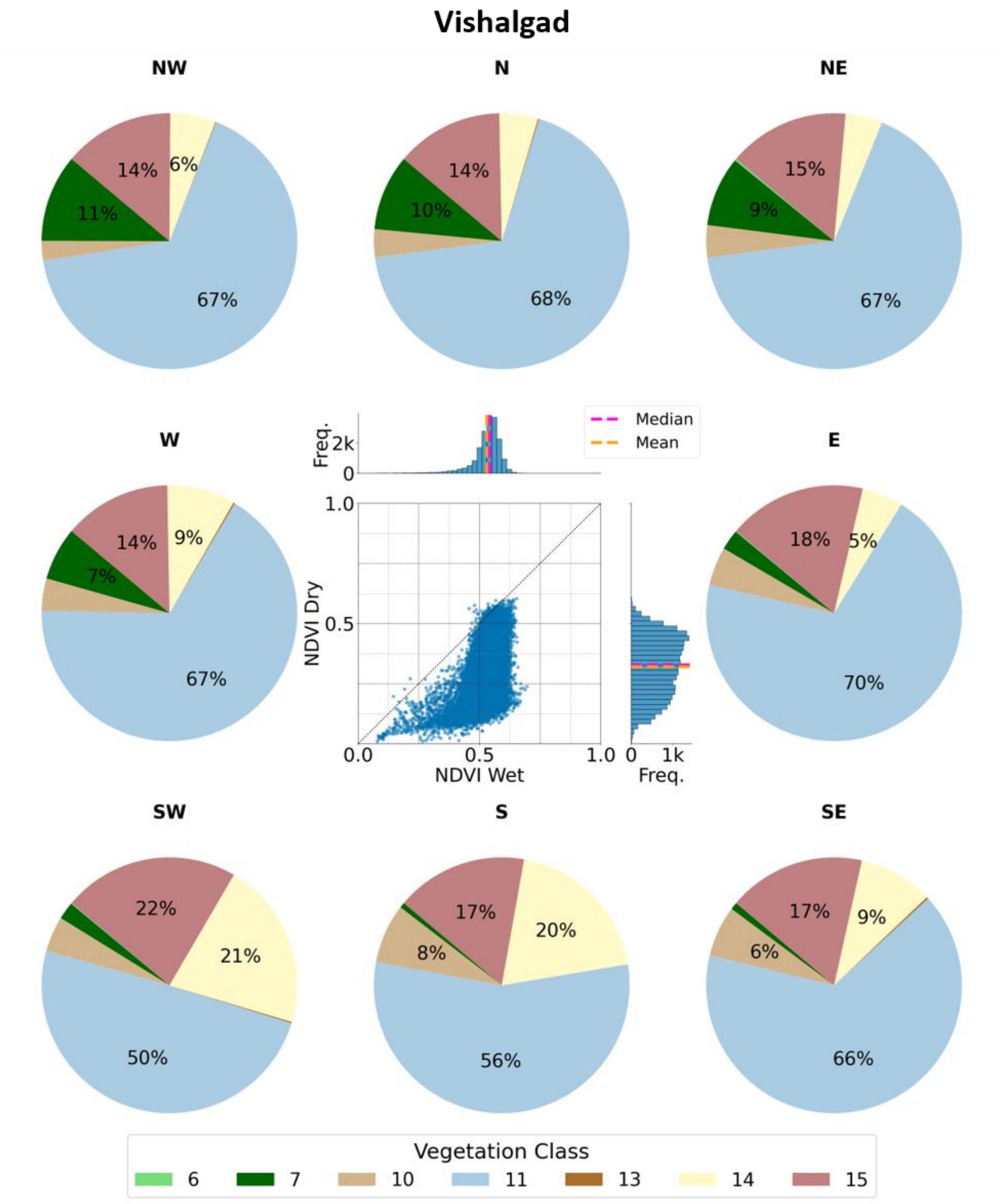


*Supplementary Figure S4. The same as Supplementary Figure S1 but for Vishalgad.*

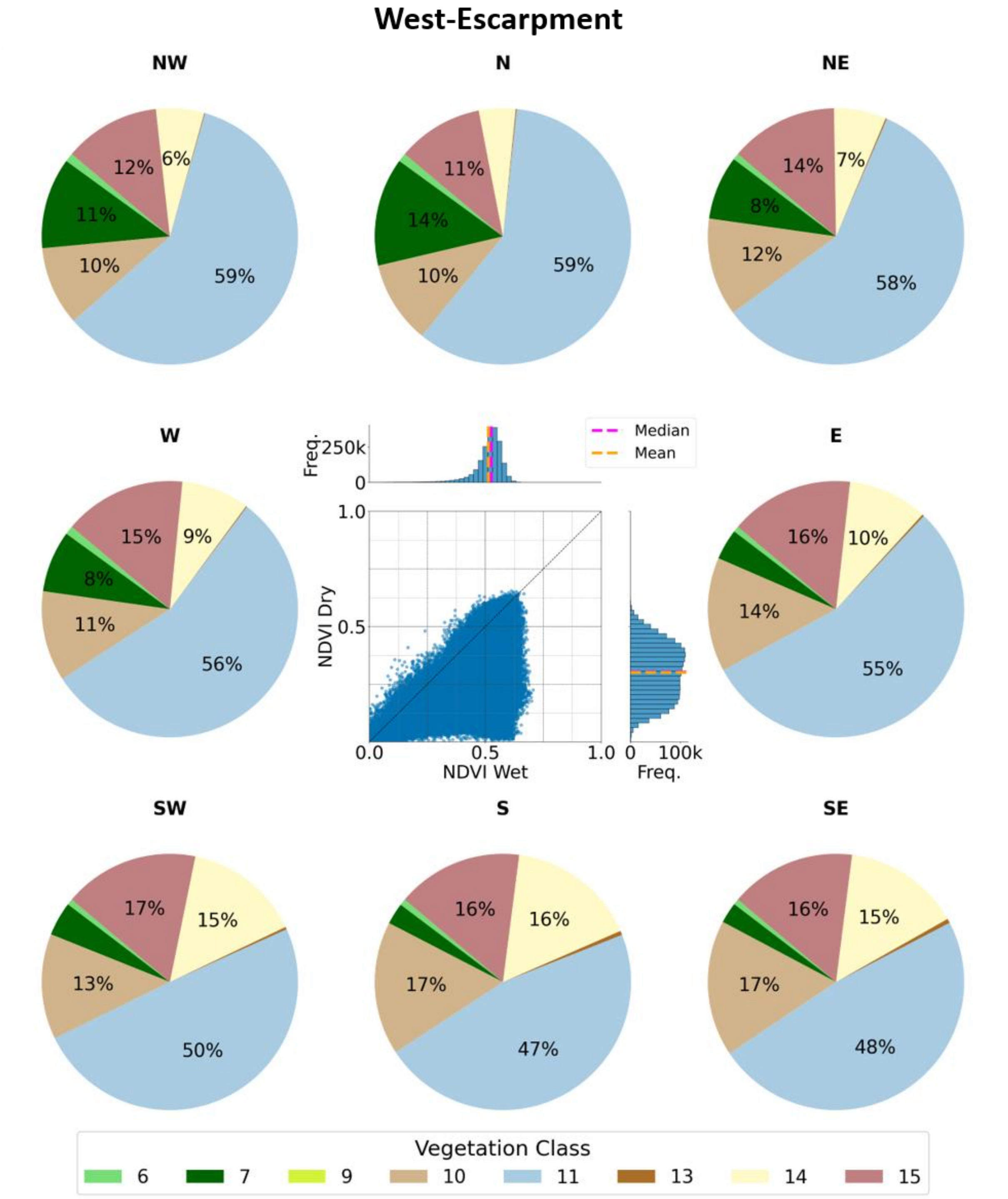


*Supplementary Figure S5. The same as Supplementary Figure S1 but for West of the WG Escarpment.*

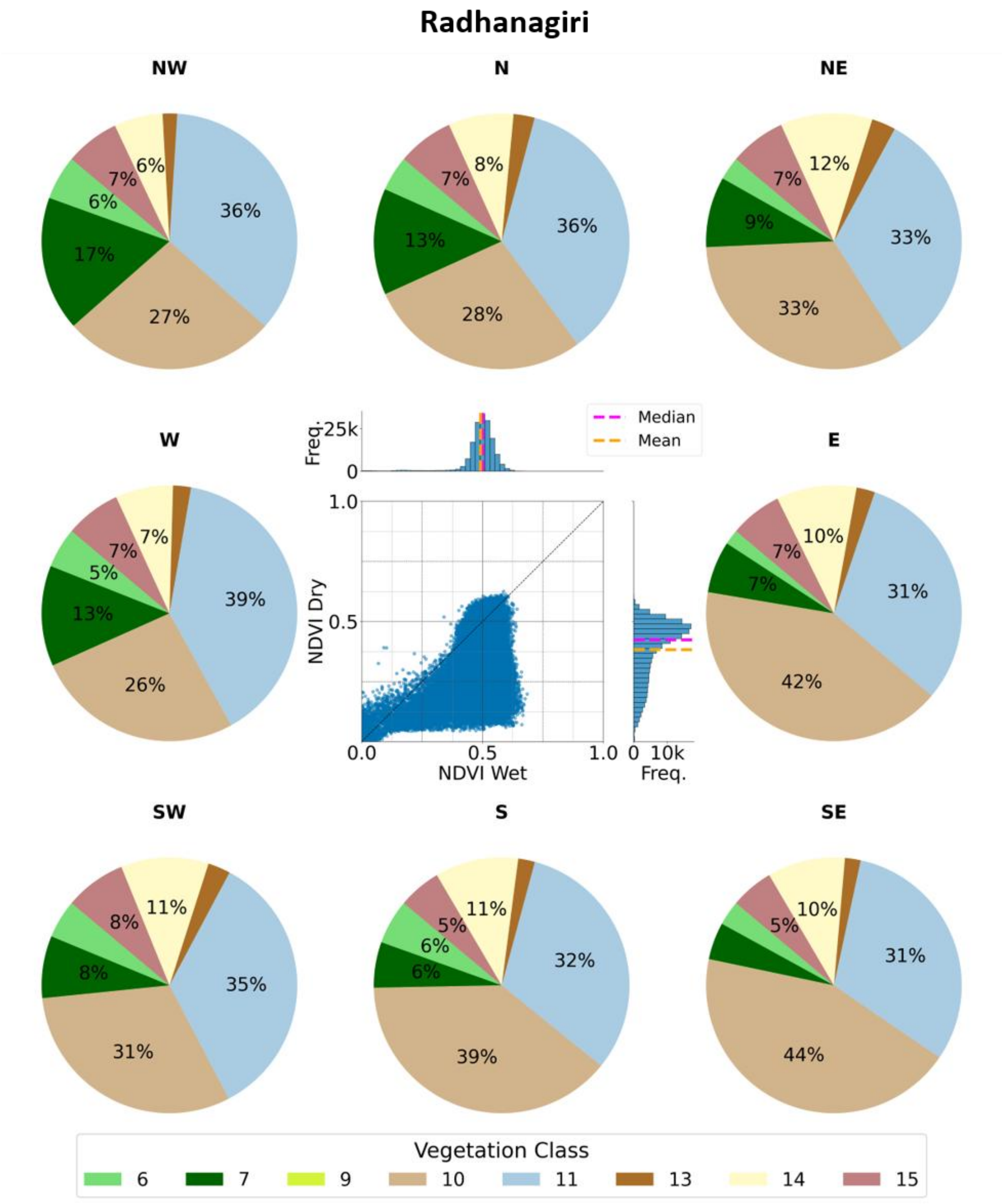


*Supplementary Figure S6. The same as Supplementary Figure S1 but for Radhanagiri.*

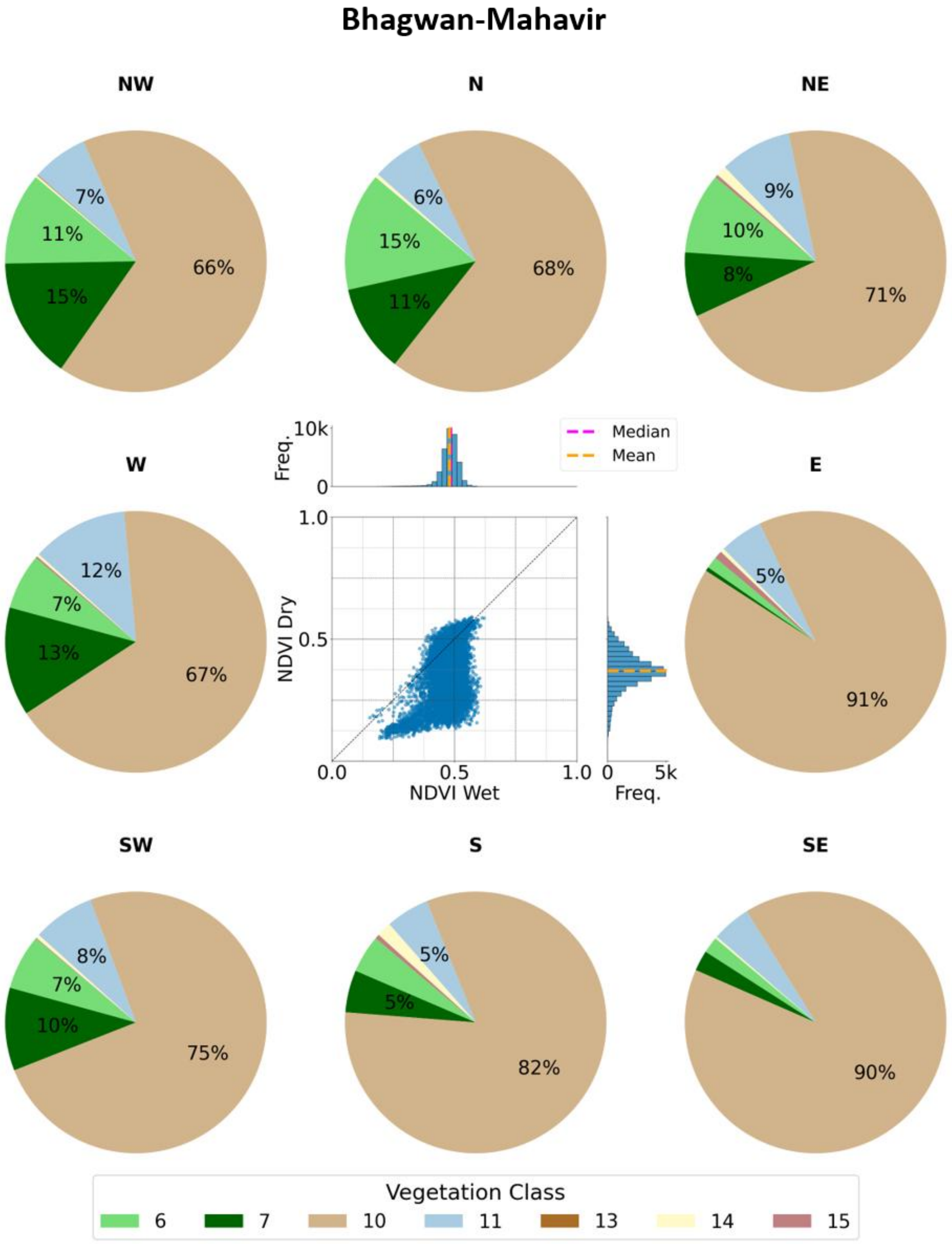


*Supplementary Figure S7. The same as Supplementary Figure S1 but for Bhagwan-Mahavir.*

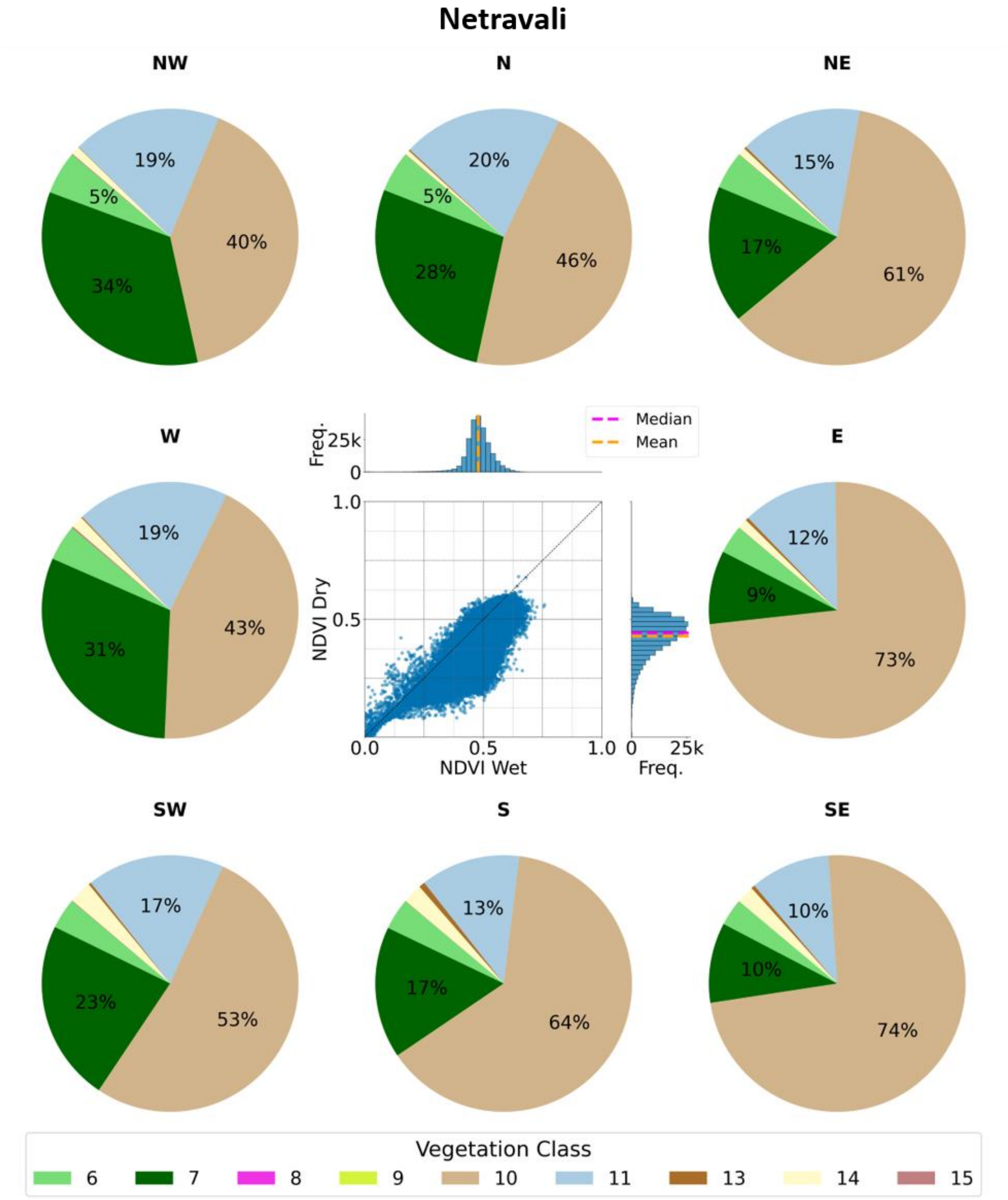


*Supplementary Figure S8: The same as Supplementary Figure S1 but for Netravali.*

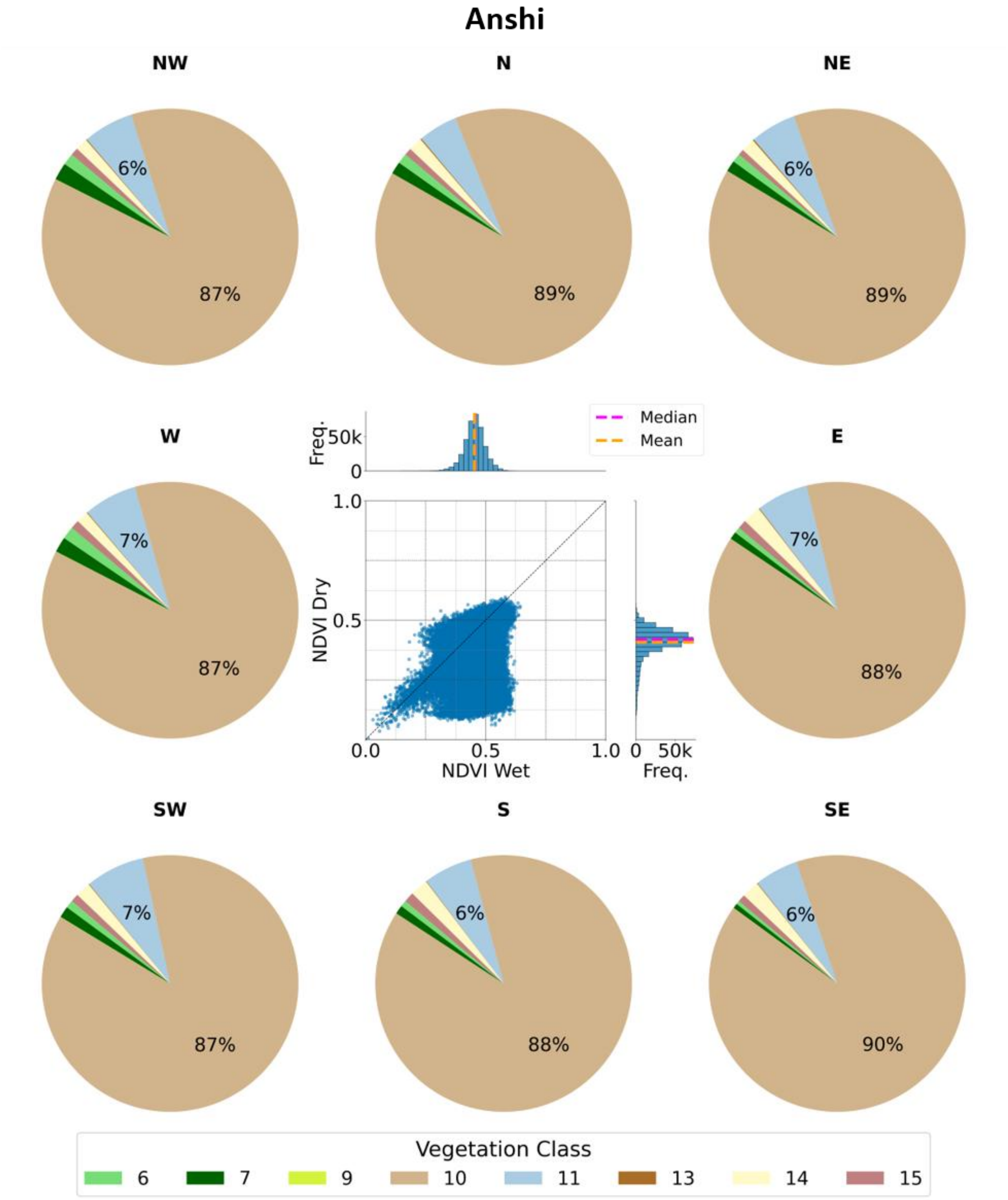


*Supplementary Figure S9. The same as Supplementary Figure S1 but for Anshi.*

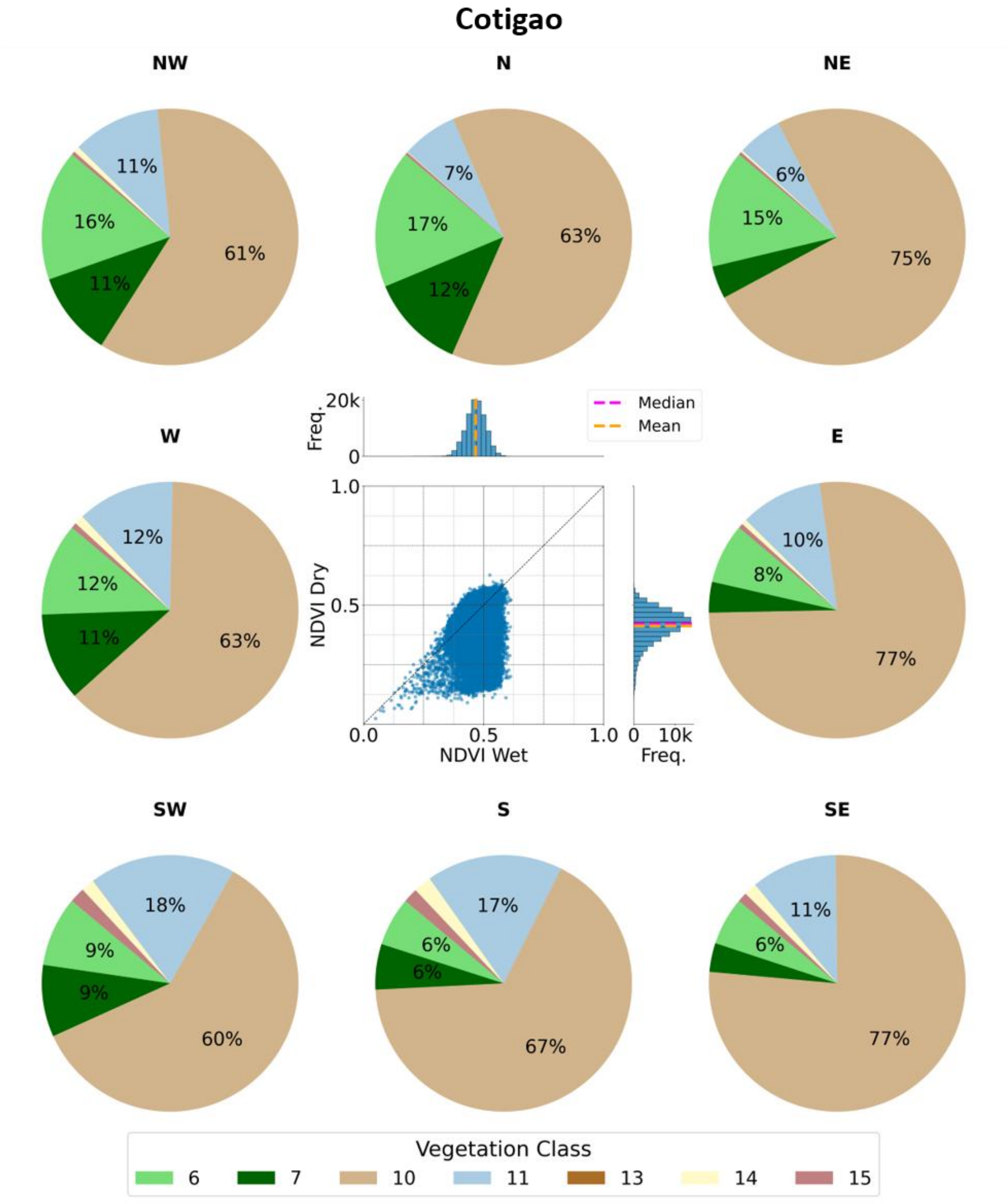


*Supplementary Figure S10. The same as Supplementary Figure S1 but for Cotigao.*

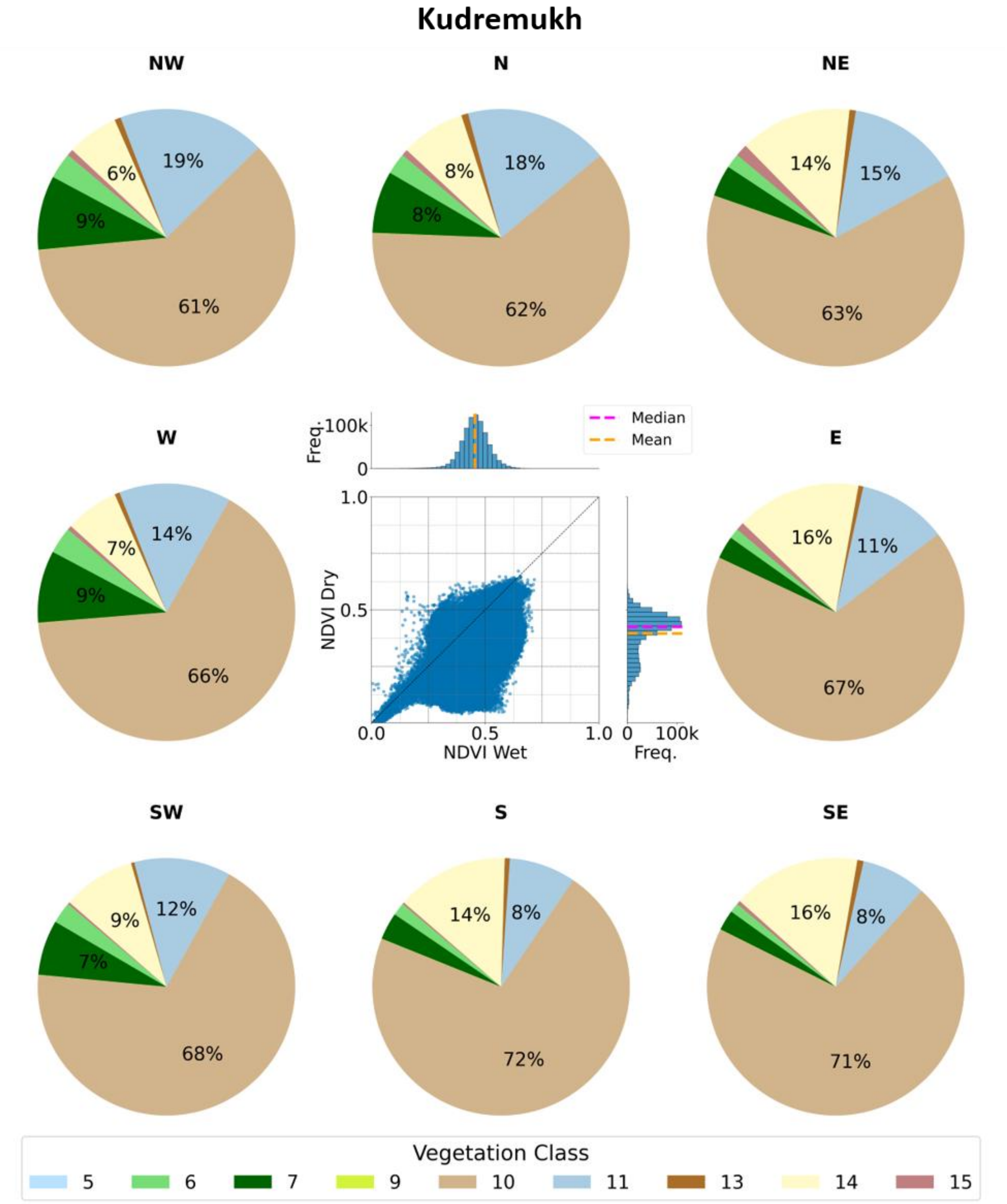


*Supplementary Figure S11. The same as Supplementary Figure S1 but for Kudremukh.*

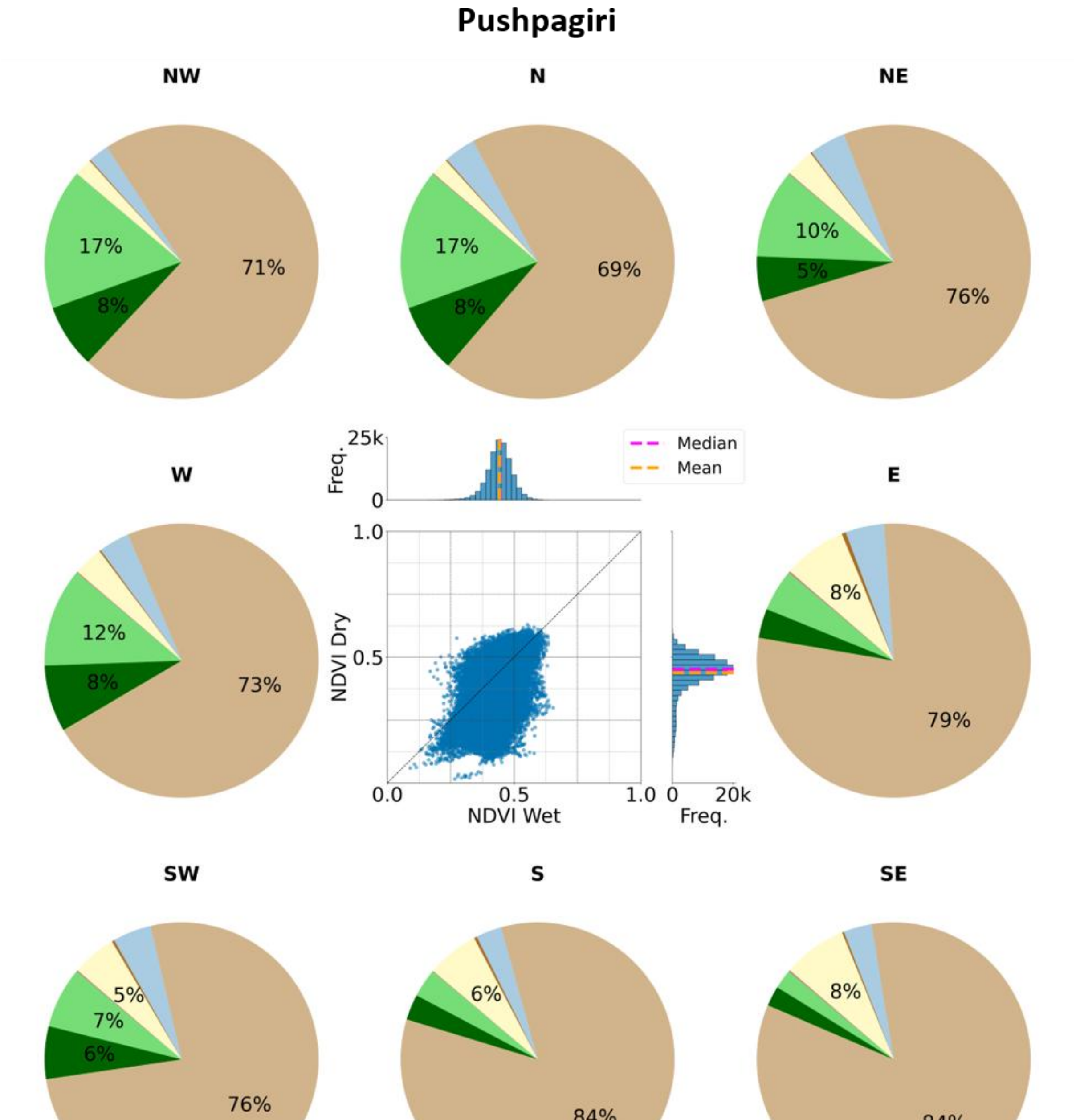


*Supplementary Figure S12. The same as Supplementary Figure S1 but for Pushpagiri.*

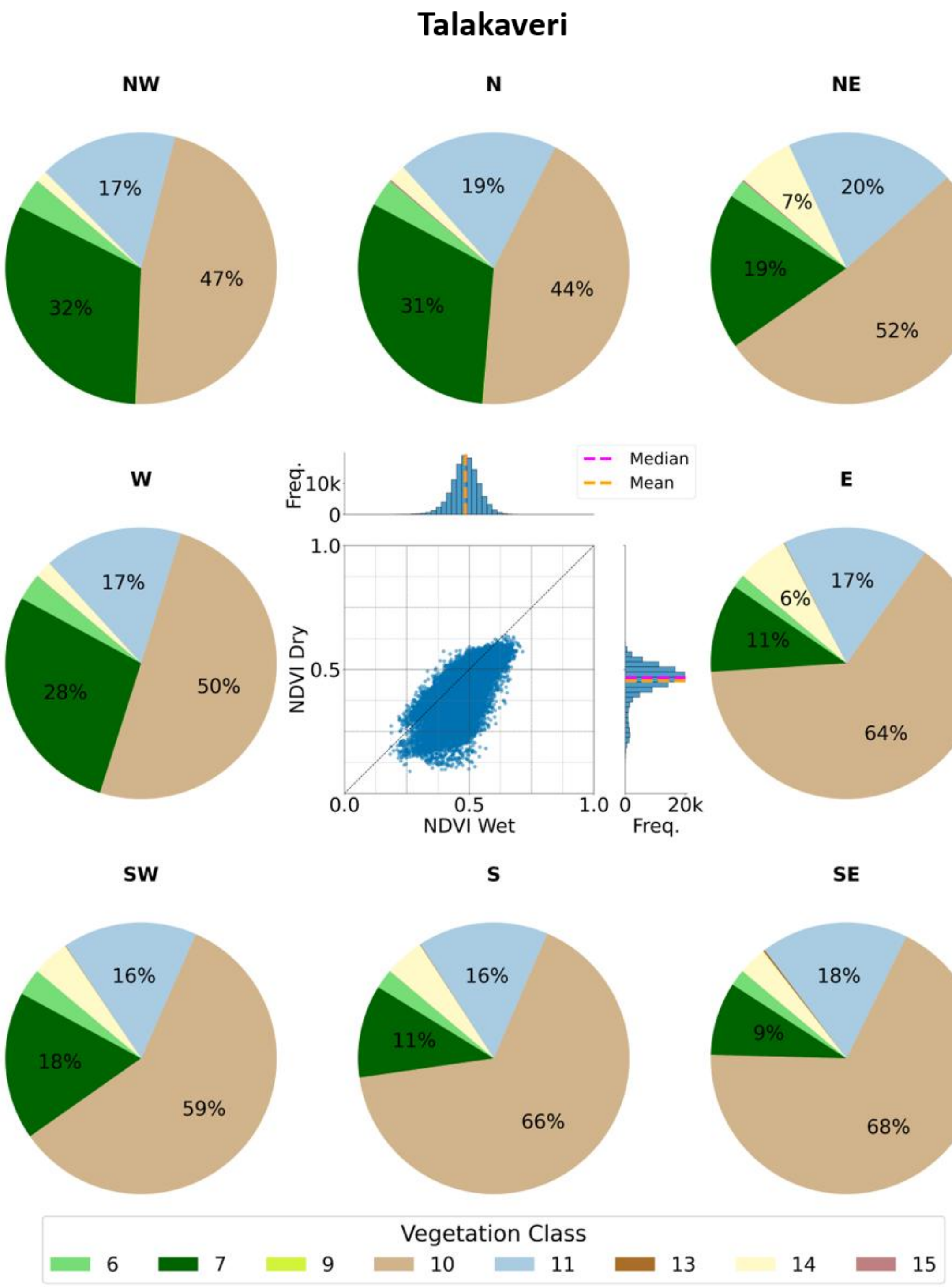


*Supplementary Figure S13. The same as Supplementary Figure S1 but for Talakaveri.*

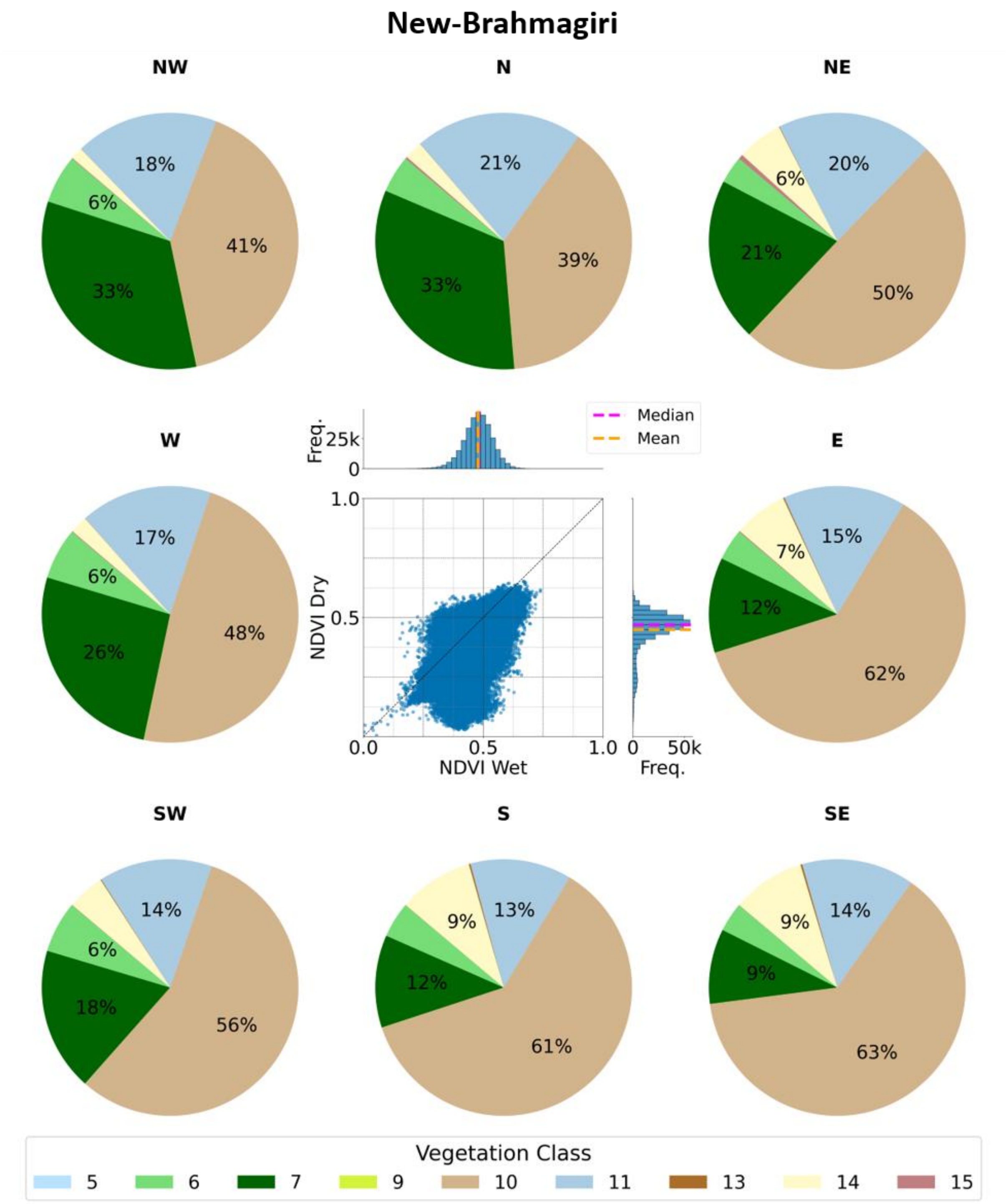


*Supplementary Figure S14. The same as Supplementary Figure S1 but for New-Brahmagiri.*

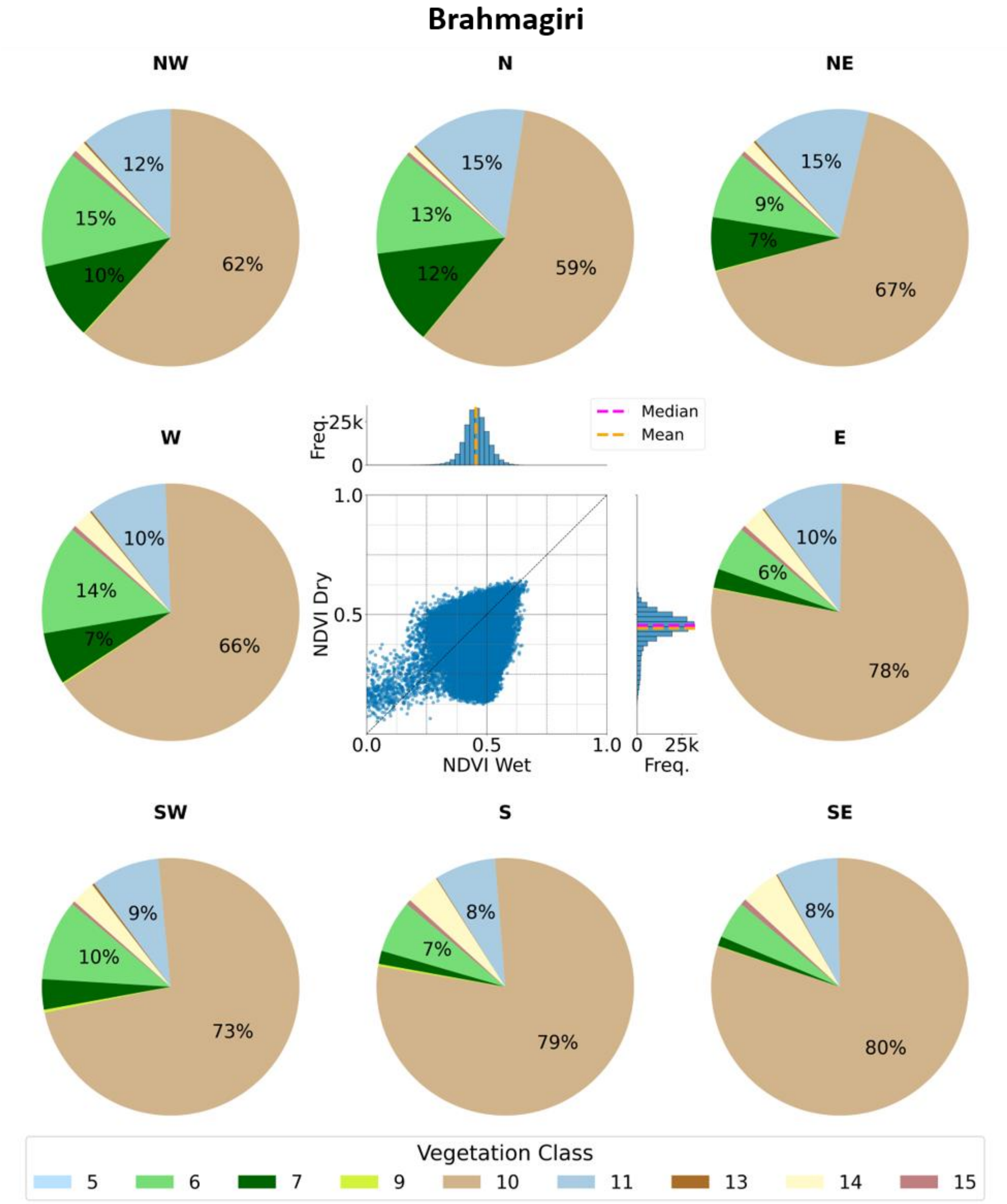


96
97
98 *Supplementary Figure S15. The same as Supplementary Figure S1 but for Brahmagiri.*
99

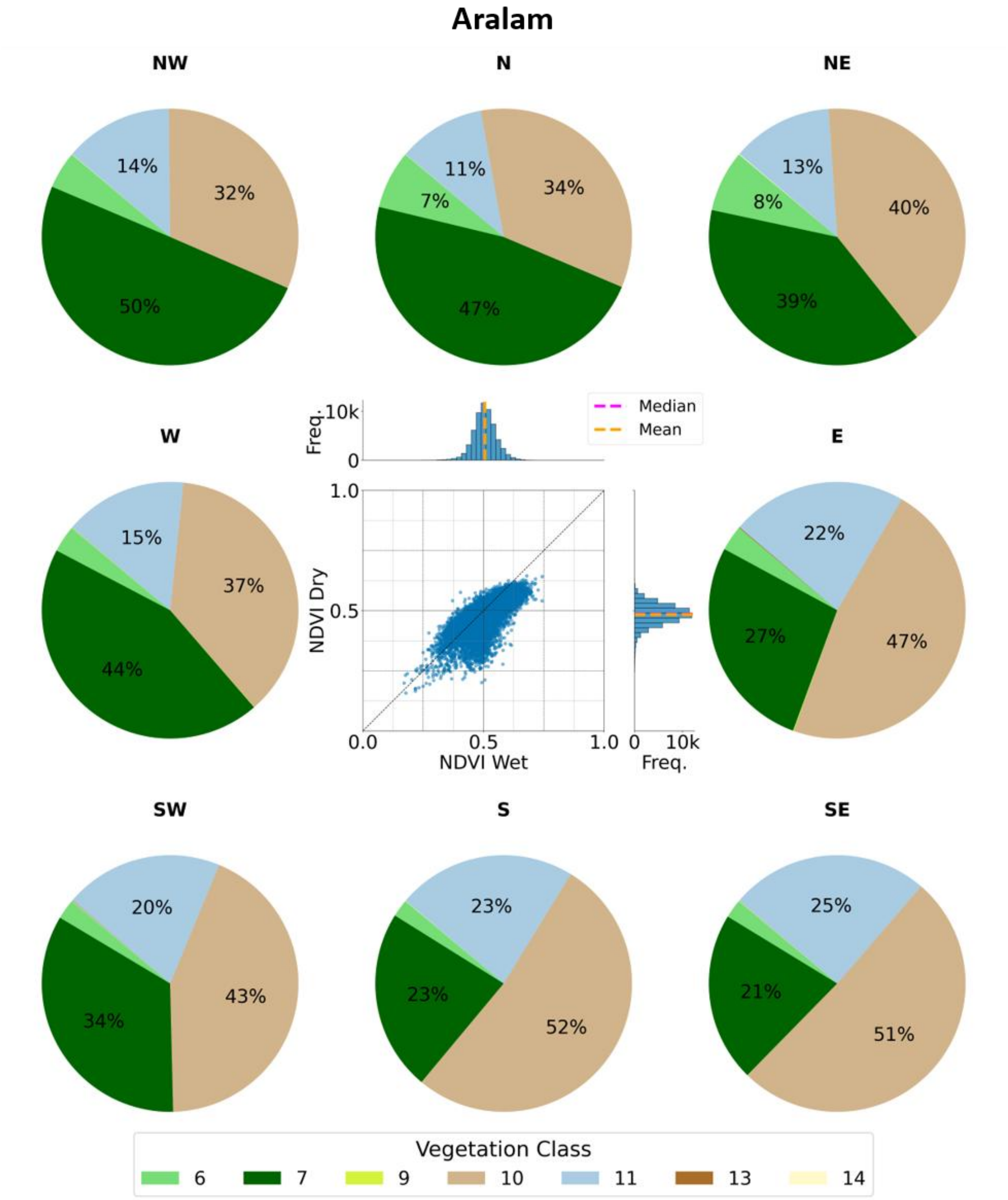


*Supplementary Figure S16. The same as Supplementary Figure S1 but for Aralam.*

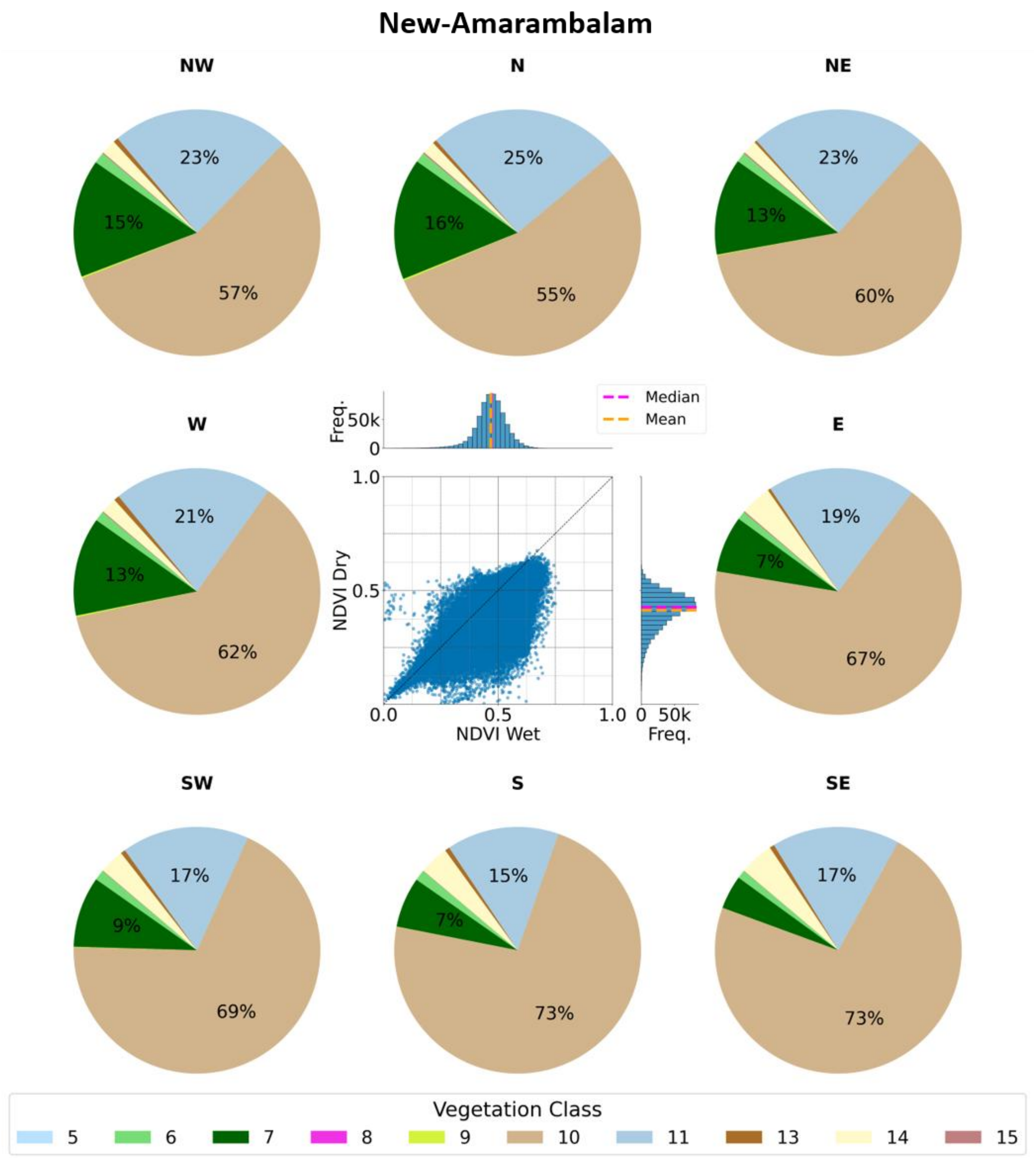


*Supplementary Figure S17. The same as Supplementary Figure S1 but for New-Amarambalam.*

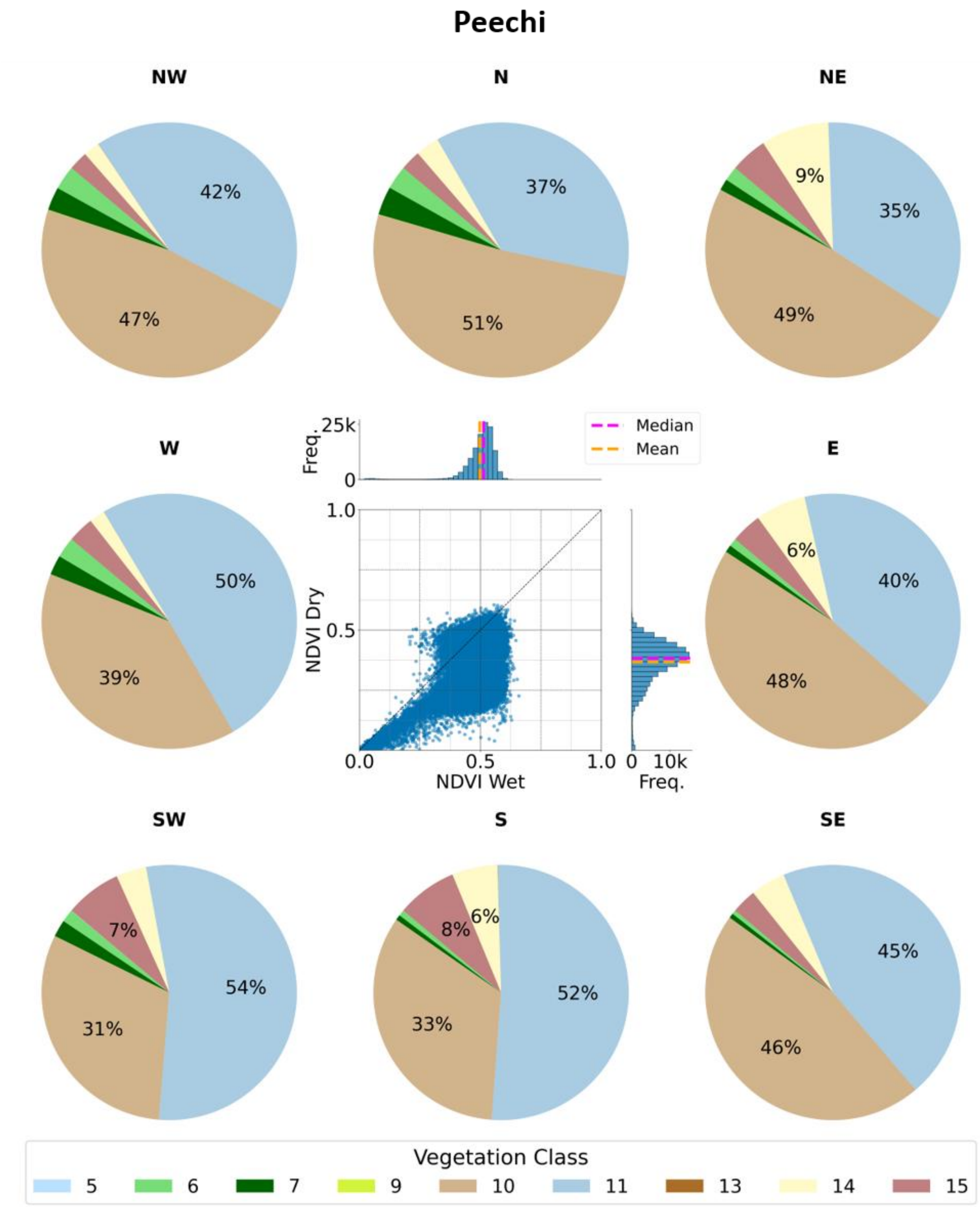


*Supplementary Figure S18. The same as Supplementary Figure S1 but for Peechi.*

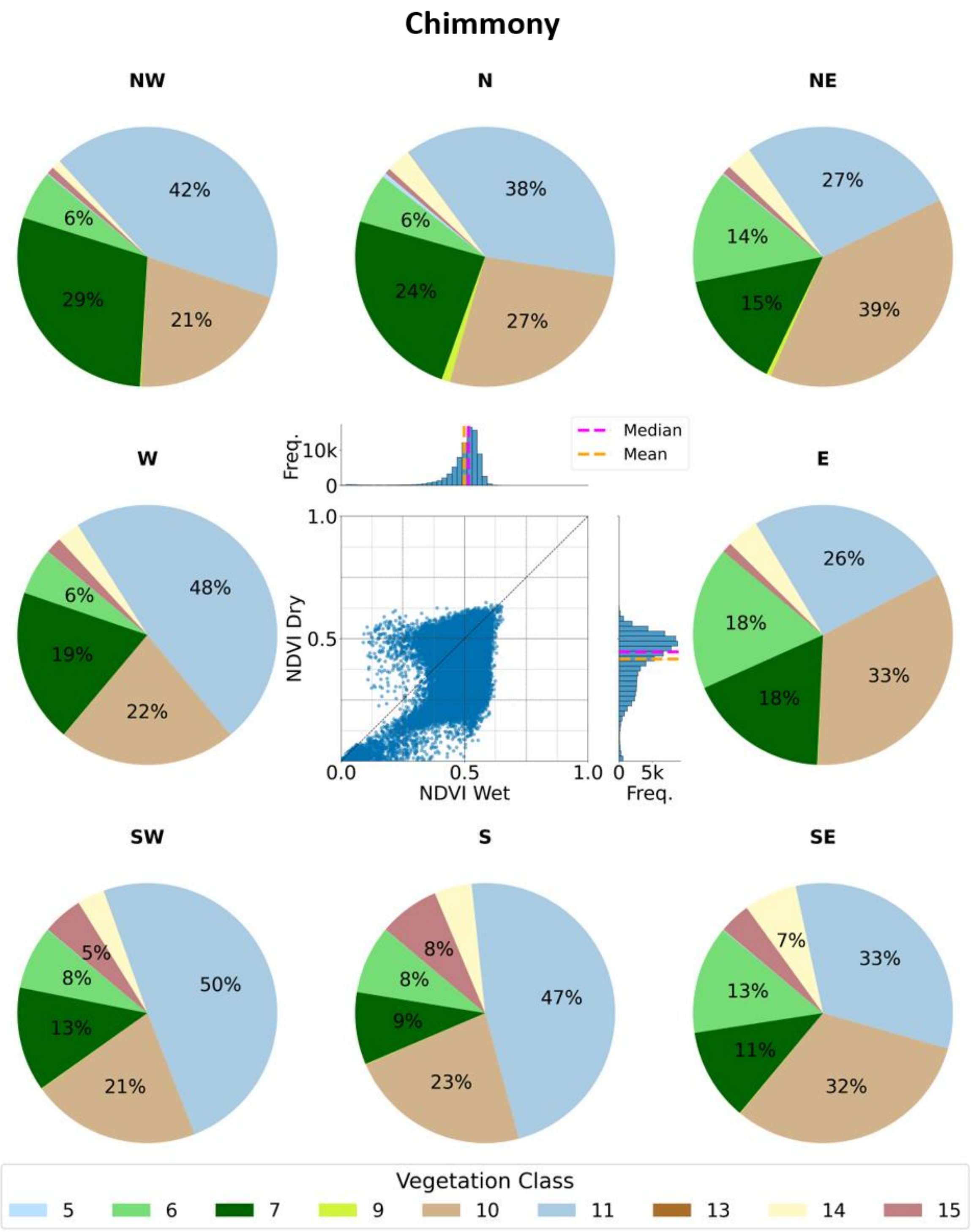


*Supplementary Figure S19. The same as Supplementary Figure S1 but for Chimmony.*

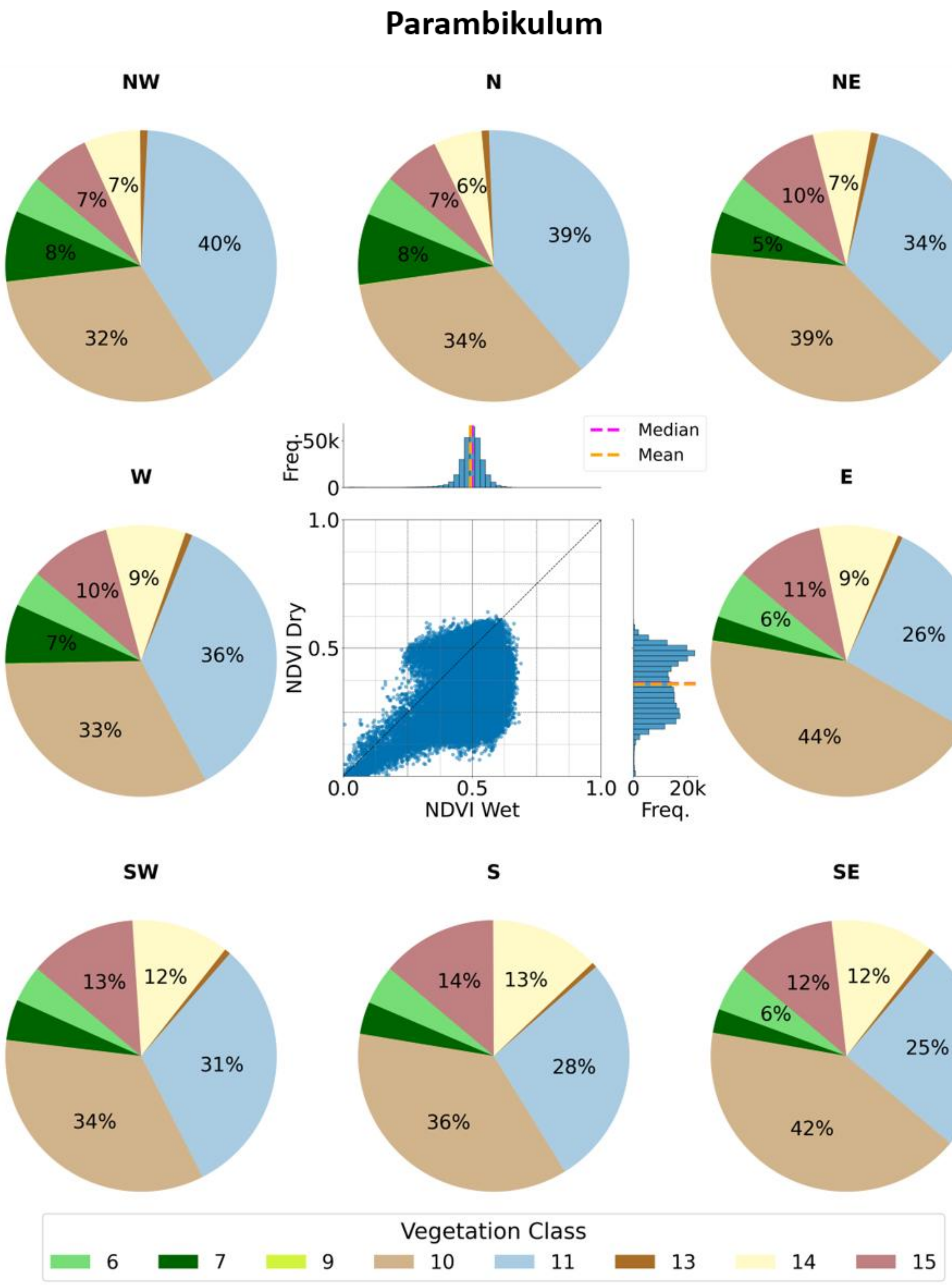


*Supplementary Figure S20. The same as Supplementary Figure S1 but for Parambikulum.*

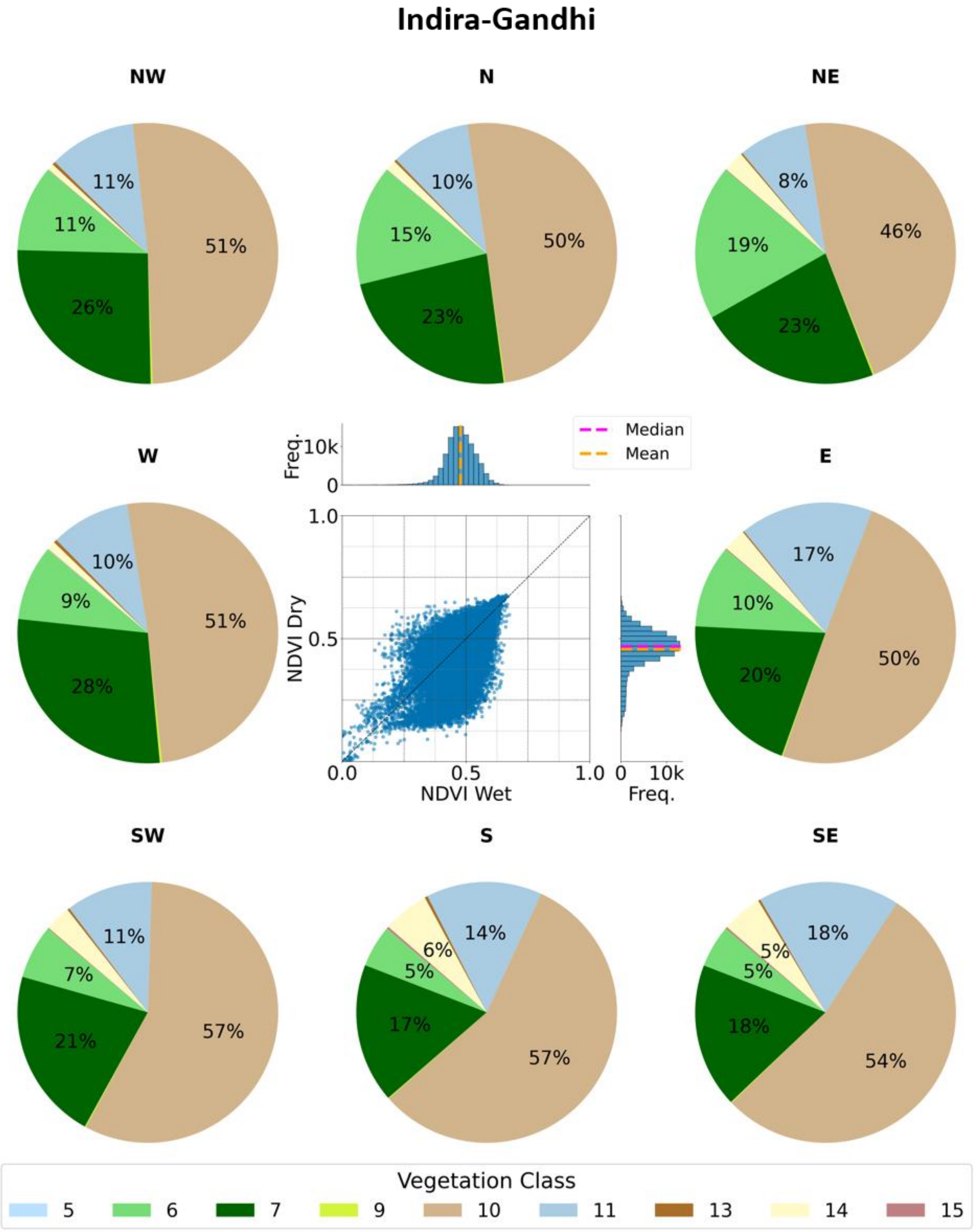


*Supplementary Figure S21. The same as Supplementary Figure S1 but for Indira-Gandhi.*

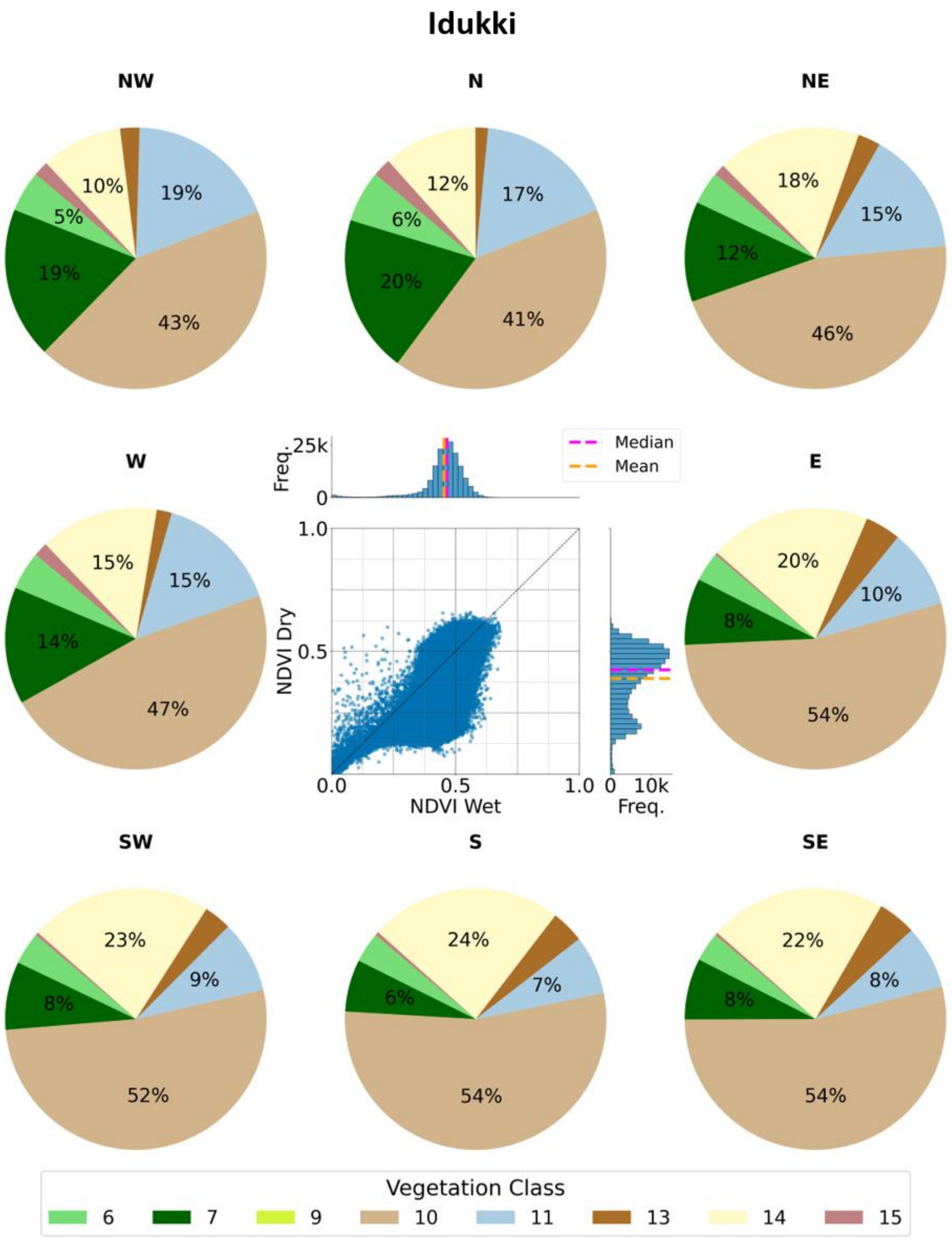


*Supplementary Figure S22. The same as Supplementary Figure S1 but for Idukki.*

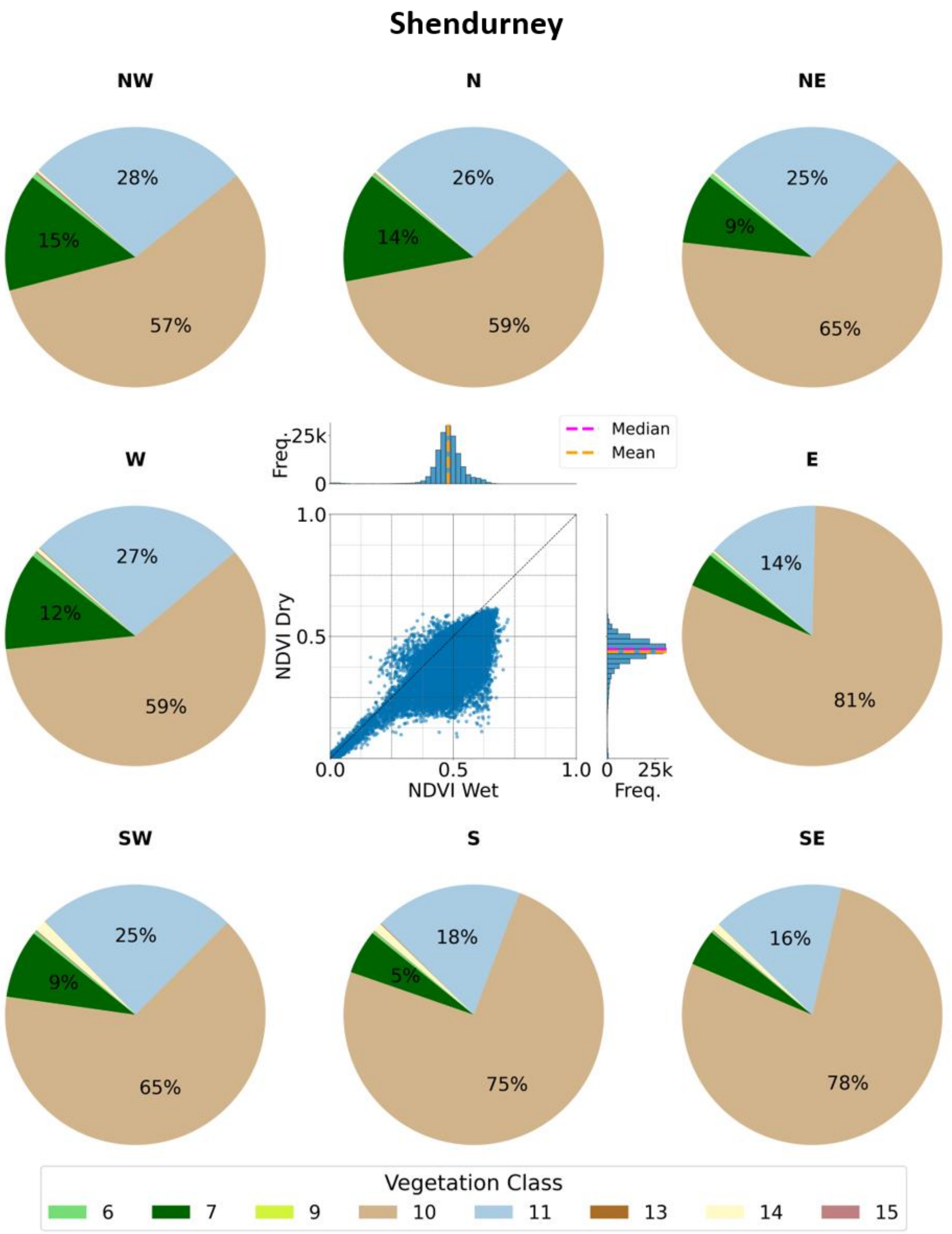


*Supplementary Figure S23. The same as Supplementary Figure S1 but for Shendurney.*

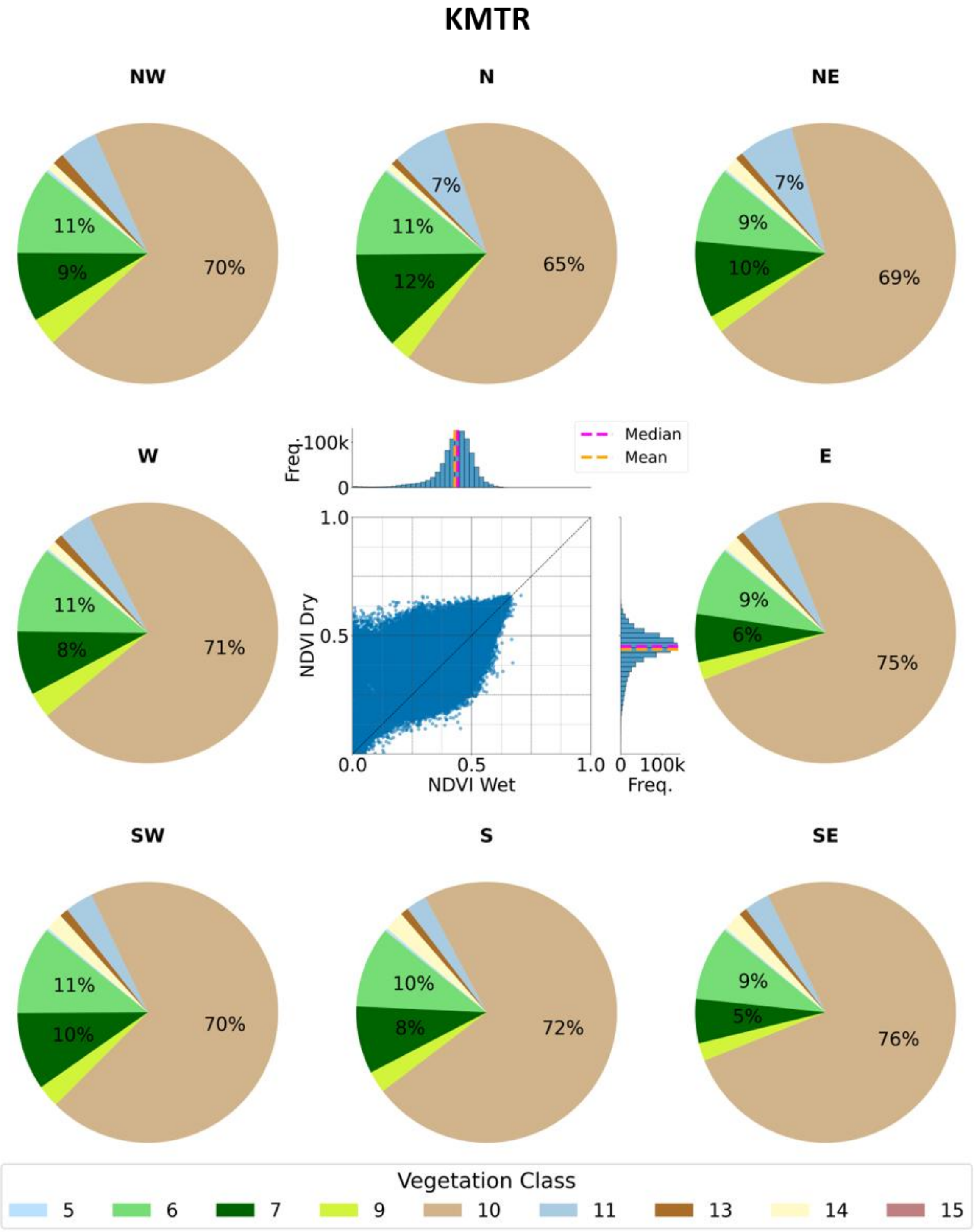


*Supplementary Figure S24. The same as Supplementary Figure S1 but for Kalakad Mundanthurai Tiger Reserve (KMTR).*

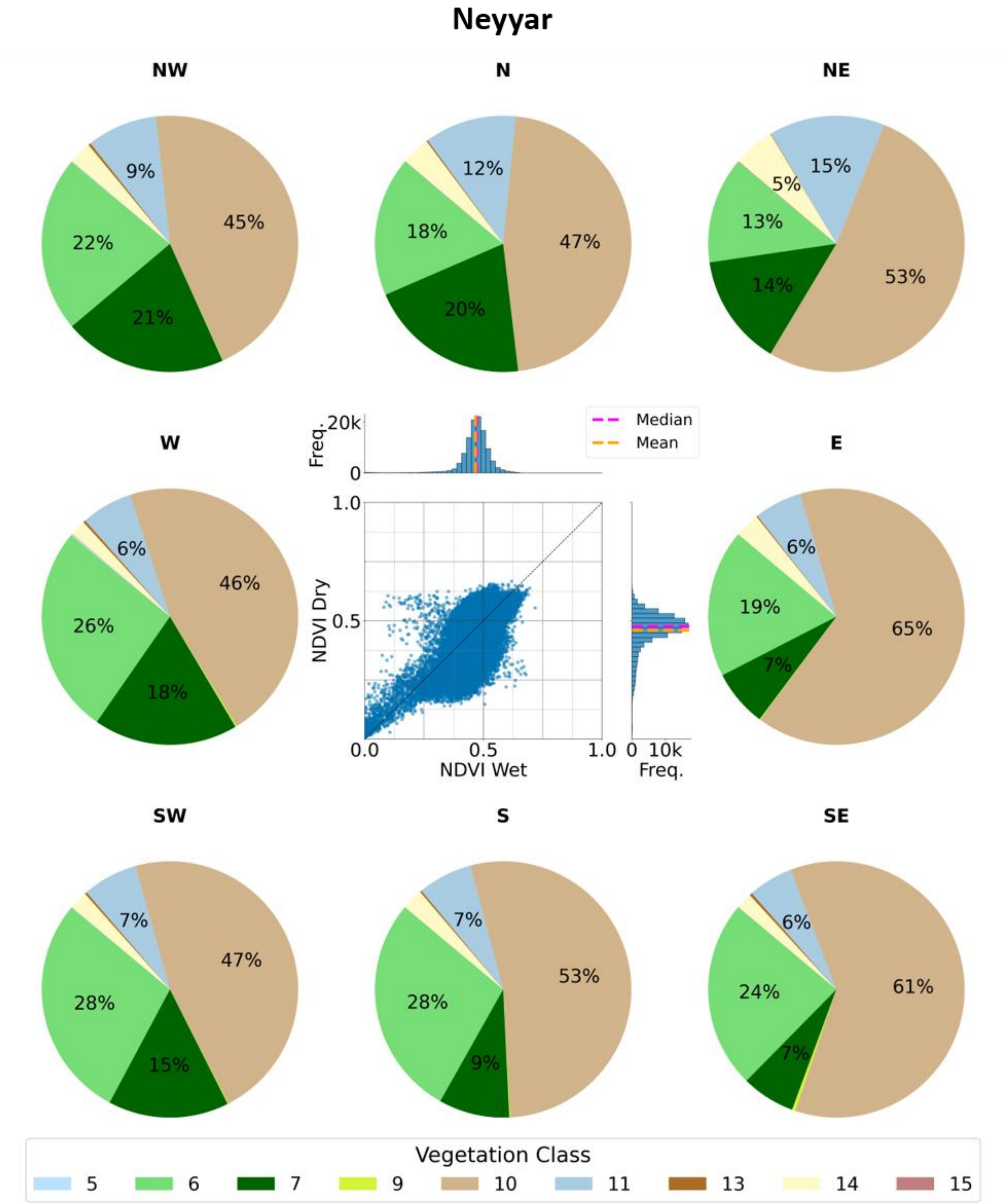


*Supplementary Figure S25. The same as Supplementary Figure S1 but for Neyyar.*
